# Inhibition of the YAP-MMB interaction and targeting NEK2 as potential therapeutic strategies for YAP-driven cancers

**DOI:** 10.1101/2023.07.27.550917

**Authors:** Marco Jessen, Dörthe Gertzmann, Franziska Liss, Franziska Zenk, Laura Bähner, Victoria Schöffler, Carsten P. Ade, Björn von Eyss, Stefan Gaubatz

## Abstract

YAP activation in cancer is linked to poor outcomes, making it an attractive therapeutic target. Previous research focused on blocking the interaction of YAP with TEAD transcription factors. Here, we took a different approach by disrupting YAP’s binding to the transcription factor B-MYB using MY-COMP, a fragment of B-MYB containing the YAP binding domain fused to a nuclear localization signal. MY-COMP induced cell cycle defects, nuclear abnormalities, and polyploidization. In an AKT and YAP-driven liver cancer model, MY-COMP significantly reduced liver tumorigenesis, highlighting the importance of the YAP-B-MYB interaction in tumor development. MY-COMP also perturbed the cell cycle progression of YAP-dependent uveal melanoma cells but not of YAP-independent cutaneous melanoma cell lines. It counteracted YAP-dependent expression of MMB-regulated cell cycle genes, explaining the observed effects. We also identified NIMA-related kinase (NEK2) as a downstream target of YAP and B-MYB, promoting YAP-driven transformation by facilitating centrosome clustering and inhibiting multipolar mitosis.

## INTRODUCTION

YAP and TAZ, the downstream effectors of the Hippo signaling pathway, are activated in many human tumors and have been identified as potent inducers of tumor cell proliferation ^1^. The Hippo signaling pathway involves multiple kinases that ultimately phosphorylate YAP and TAZ, triggering their cytoplasmic retention and degradation via the proteasome ^2, 3^. One mechanism of YAP activation in cancer is through the loss of function mutations or deletions in genes encoding key components of the Hippo pathway, such as NF2 and LATS1/2 ^4^. Loss of function mutations or deletions in these genes can lead to reduced phosphorylation and inactivation of YAP/ TAZ, resulting in their translocation to the nucleus and subsequent transcriptional activation of genes involved in cell proliferation and survival. In addition, YAP/TAZ activation can be mediated by other signaling pathways. For example, in uveal melanoma, oncogenic mutations in G protein-coupled receptor (GPCR) stimulate the Rho GTPase and the actin cytoskeleton, which in turn inhibits Hippo pathway signaling and promotes YAP/TAZ activation ^5, 6^.

YAP/TAZ do not directly bind to DNA but function as transcriptional coactivators by interacting with TEAD transcription factors ^2, 3, 7, 8^, which recruit YAP/TAZ primarily to distal transcriptional enhancers, regulatory DNA elements that activate the expression of their target genes by the formation of chromatin loops ^9–13^. While TEAD transcription factors are primarily responsible for the recruitment of YAP, YAP also collaborates with other transcription factors and co-activators. For example, YAP-regulated enhancers often contain both TEAD and AP-1 motifs, where YAP synergizes with JUN/FOS to promote tumor cell proliferation and transformation ^10, 14, 15^. There are ongoing efforts to develop inhibitors that target the YAP-TEAD interaction, which represents a promising therapeutic strategy for cancer treatment ^16^.

In previous work we have shown that YAP binds to and cooperates with the B-MYB transcription factor, a subunit of the Myb-MuvB (MMB) complex to activate cell cycle dependent gene expression ^17, 18^. Mechanistically, by binding to distant enhancers, YAP promotes the binding of B-MYB to MuvB at the transcription start site (TSS) of its target genes. Once the YAP-B-MYB-MuvB complex is formed, it activates the expression of genes involved in the G2/M progression, mitosis and cytokinesis. Moreover, YAP increases the expression of B-MYB, which contributes to an increased rate of mitosis and hyperproliferation. Based on identification of the YAP-binding domain in B-MYB, we have generated MY-COMP (for B-<u>M</u>YB-<u>Y</u>AP <u>comp</u>etition), a fragment of B-MYB that, when overexpressed binds to YAP and interferes with the endogenous YAP-B-MYB interaction ^18^. Disrupting the YAP-B-MYB interaction in cardiomyocytes with MY-COMP strongly inhibits their proliferation.

Here we have investigated the significance of the YAP-B-MYB interaction for YAP-dependent tumorigenesis. We find that MY-COMP antagonizes the YAP-dependent expression of MMB-target genes but has weak effects on gene expression in normal pre-malignant MCF10A cells. *In vivo*, MY-COMP inhibits tumorigenesis in a mouse model of liver cancer driven by activated AKT and YAP. Furthermore, YAP-dependent uveal melanoma cells, but not YAP-independent cutaneous melanoma cell lines are sensitive to the blockage of the YAP-B-MYB interaction by MY-COMP. We also identified NIMA-related kinase (NEK2) as a candidate target downstream of YAP contributing to the transformation of YAP-dependent uveal melanoma cells. Targeting selected YAP-MMB regulated genes such as NEK2, or suppression of YAP-regulated cell cycle genes by inhibiting the WW-domains of YAP could provide a novel strategy to antagonize the pro-tumorigenic functions of YAP.

## RESULTS

In our previous study, we demonstrated that the Hippo coactivator YAP directly interacts with the B-MYB transcription factor, a subunit of the Myb-MuvB complex ^18^. Based on the identification of the YAP-binding domain in B-MYB, we generated MY-COMP (for B-<u>M</u>YB-<u>Y</u>AP <u>comp</u>etition), consisting of amino acids 2-241 of B-MYB fused to a nuclear localization signal (Figure 1A). We have previously demonstrated that MY-COMP interferes with the endogenous YAP-B-MYB interaction in murine cardiomyocytes ^18^. In this study we addressed the consequence of MY-COMP expression in human cancer cells and in a mouse tumor model driven by oncogenic YAP. To do so, we first created HeLa cells stably expressing doxycycline inducible MY-COMP linked to GFP via a T2A site (Figure 1B). To test whether MY-COMP is able to inhibit binding of B-MYB to YAP, we performed co-immunoprecipitation experiments with Flag-tagged YAP and HA-tagged B-MYB. Cells only expressing flag-tagged YAP served as control. Without doxycycline, HA-B-MYB robustly co-precipitated with flag-YAP (Figure 1C). After induction of MY-COMP, the interaction between Flag-YAP and HA-B-MYB was reduced. As a control, deletion of the WW-domains in YAP results in complete loss of the YAP-B-MYB interaction, as expected. To test how MY-COMP effects cell cycle progression, we next performed FACS analysis after 4 days of doxycycline treatment. Cells expressing MY-COMP were identified by their green fluorescence. Upon expression of MY-COMP, the fraction of polyploid cells strongly increased from 12.5% to more than 47%. In parallel, the fraction of cells in G1 phase decreased after MY-COMP was expressed (Figure 1D,E). Importantly, there were no changes in the cell cycle phases upon expression of a shorter version of MY-COMP(2-79) that does not interact with YAP, confirming the specificity of MY-COMP (Figure 1B,E,F). To analyze the defects in cell division by MY-COMP in more detail, we expressed MY-COMP and a set of MY-COMP mutants in HeLa cells. In addition to the shortened MY-COMP(2-79) construct introduced above, we introduced a mutation into the DNA-binding domain of MY-COMP that is expected to prevent the interaction of B-MYB with DNA ^19^. Finally, because we have previously shown that binding of YAP to B-MYB is mediated by a PPXY motif in B-MYB, we used a MY-COMP mutant in which the PPXY motif was deleted.

**Figure 1:**
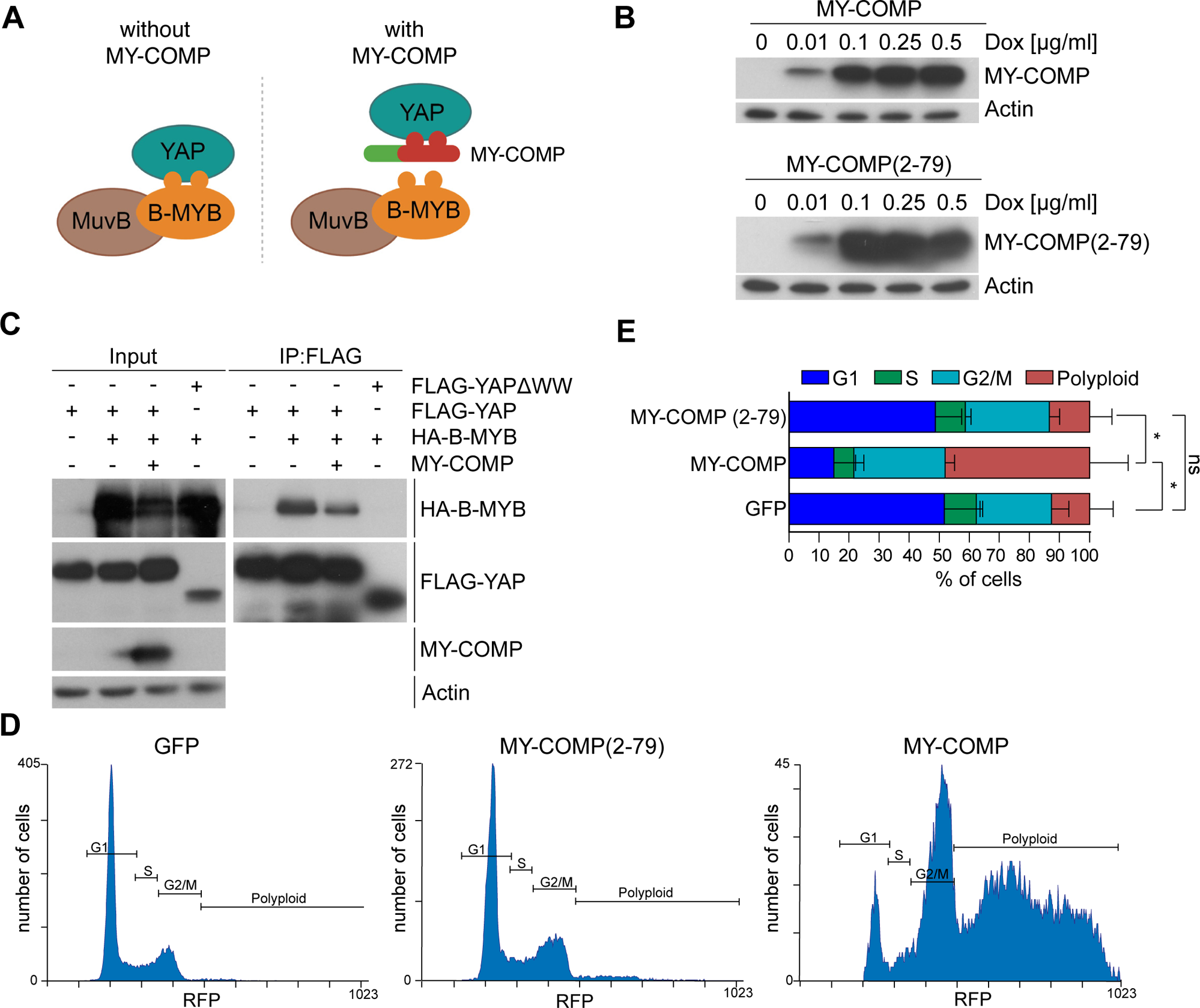
MY-COMP disrupts the interaction of YAP with B-MYB and leads to polyploidy. A) Scheme for the disruption of the YAP-B-MYB interaction by MY-COMP. B) HeLa cells were stably transfected with expression constructs for inducible MY-COMP or MY-COMP (2-79). Cell lines were treated with the indicated dose of doxycycline and the expression of MY-COMP was analyzed by immunoblotting with an anti-HA antibody. Actin served as a loading control. C) HeLa cells were transfected with the indicated expression constructs. Flag-YAP was immunoprecipitated and bound HA-B-MYB was detected by immunoblotting. When MY-COMP was co-transfected, the interaction between flag-YAP and HA-B-MYB was decreased. Input: 3 % of the lysate. D) The expression of GFP or of the indicated MY-COMP constructs coupled to GFP through a T2A site was induced by doxycycline. FACS was performed to determine the percentage of in different phases of the cell cycle. GFP-positive cells (expressing MY-COMP) were analyzed. E) Quantification of the results shown in D. Error bars indicate SDs. n=3 independent replicates. Student’s t-test. *=p<0.05, ns= not significant.

Immunoblotting confirmed that mutant MY-COMP constructs were expressed at comparable levels (Figure 2A). Next, we performed immunofluorescence staining where MY-COMP expressing cells were identified by staining with a HA antibody (Figure 2B). Strikingly, expression of MY-COMP resulted in a pronounced increase in bi- and multinucleated cells as well as in cells with micronuclei (Figure 2B,C). MY-COMP(2-79) showed no such effect. The effect of MY-COMP is not due to interference with the DNA-binding of B-MYB, because MY-COMP with the mutation in the DNA-binding domain showed the same phenotype ^19^. This is expected, because DNA-binding of MMB is not mediated by B-MYB, but by the LIN54 subunit of MuvB ^20, 21^. Deletion of the PPXY motif in MY-COMP abolished the effect on bi- and multinucleation and decreased micronuclei formation, indicating that the ability to induce cell division errors correlates with the ability to disrupt the YAP-B-MYB interaction. Because MY-COMP constructs were expressed at similar levels, the failure of the mutants to cause a phenotype cannot be due to lower expression of these constructs. We did not find any changes in YAP expression levels and YAP S217 phosphorylation in cells expressing MY-COMP (Figure 2A). Phosphorylation of S127 was used as a proxy for YAP activity as it is a key residue for phosphorylation by the Hippo kinases LATS1/2. Thus, MY-COMP does not influence the expression of YAP and B-MYB or the activity of YAP. Taken together, these experiments indicate that the disruption of the YAP-B-MYB interaction results in errors in mitotic progression in human cells.

**Figure 2:**
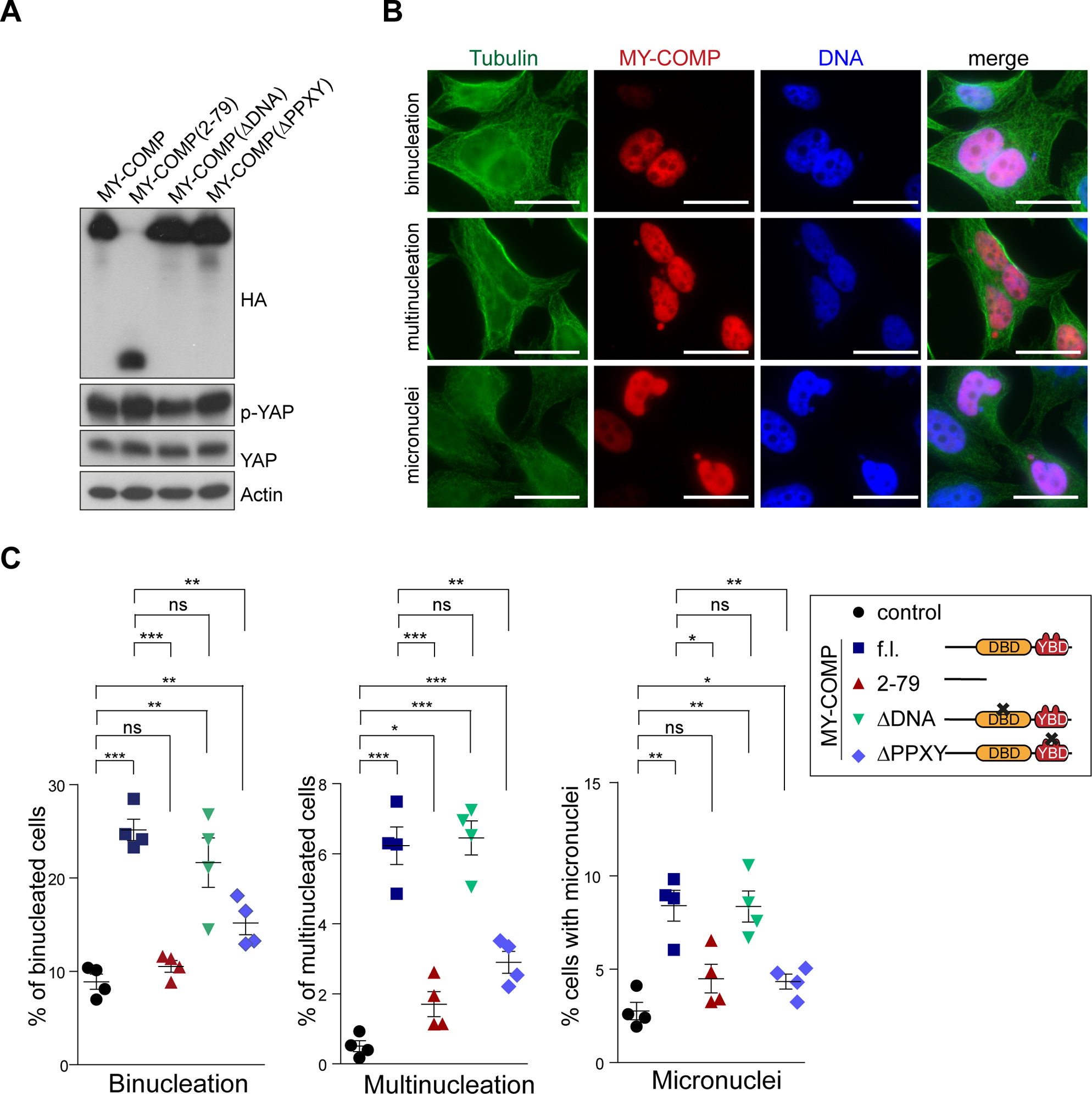
MY-COMP leads to errors in cell division. A) Expression of the indicated MY-COMP constructs was verified by immunoblotting with an HA-antibody. MY-COMP constructs did not affect the expression of YAP and YAP phosphorylated on S127 (p-YAP). Actin served as a control. B) Microphotographs showing examples of the different phenotypes following expression of MY-COMP. Bar: 25 μm. C) Quantification of the fraction of binucleated and multinucleated cells and cells with micronuclei following expression of the indicated MY-COMP constructs. N=4 biological replicates. Error bars indicate SEMs. Student’s t-test. *=p<0.05, **=p<0.01, ***=p<0.001, ns= not significant.

### MY-COMP inhibits YAP mediated transcription

Next, we investigated whether MY-COMP can suppress YAP-mediated transcription of MMB target genes. To address this question, we used untransformed, pre-malignant epithelial MCF10A cells. MCF10A cells express very low levels of endogenous YAP, which is primarily cytoplasmic ^22^. We first established MCF10A cells stably expressing ER-YAP2SA. YAP2SA is a constitutively active YAP mutant in which two key LATS1/2 phosphorylation sites are mutated from S to A. YAP2SA was fused to the hormone-binding domain of the estrogen receptor, which can be activated by the addition of 4-hydroxytamoxifen (4-OHT) to the culture medium. Treatment of ER-YAP2SA cells with 4-OHT resulted in stabilization of ER-YAP2SA and translocation to the nucleus as determined by cellular fractionation and immunoblotting (Supplemental Figure S1A). Importantly, addition of 4-OHT rapidly induced the mRNA expression of known YAP target genes (Supplemental Figure S1B). Induction of TOP2A and CDC20 after ER-YAP2SA activation with 4-OHT was also confirmed on protein level by immunoblotting (Supplemental Figure S1B).

We next introduced doxycycline-inducible MY-COMP into ER-YAP2SA cells. We confirmed doxycycline-mediated nuclear expression of MY-COMP by immunostaining and immunoblotting using an antibody directed against the HA-tag (Supplemental Figure S1D,E). Importantly, MY-COMP interfered with YAP-mediated expression of CDC20 and TOP2A, but not with expression of ER-YAP2SA itself (Supplemental Figure S1E). Notably, *CYR61 mRNA* expression was not affected by MY-COMP, consistent with the notion that this gene is regulated independently from the YAP-MMB interaction by direct binding of YAP to the promoter of this gene (Supplemental Figure S1F).

To evaluate the global impact of MY-COMP on the YAP-mediated transcriptional program, we next performed RNA-sequencing and compared the effect of YAP2SA in presence and absence of MY-COMP. We found that MY-COMP interfered with both activation as well as repression by YAP2SA (Figure 3B). Overall, MY-COMP lead to an approximately twofold reduction in YAP2SA-dependent gene expression (Figure 3C). Importantly, GSE analysis of the RNA-seq data showed that MY-COMP does not block all transcription equally, but selectively acts on genes related to cell cycle and mitosis, consistent with the requirement of the YAP-B-MYB interaction for mitotic gene expression (Figure 3D) ^17, 18^. We confirmed these results by plotting specific gene sets, which showed that MY-COMP prevented the induction of MMB-target genes and E2F-targets, while genes directly regulated by YAP binding within 1 kb of the transcriptional start site (TSS) (“YAP-bound”) remained unaffected (Figure 3E). Furthermore, we observed that MY-COMP had a weaker impact on MMB-target gene expression in the absence of ER-YAP2SA induction compared to its effects after YAP2SA-induction (Figure 3F). Volcano plots further illustrate that ER-YAP2SA upregulated MMB-target genes, which was impaired by MY-COMP (Figure 3G). To validate our findings, we randomly selected several top hits of YAP-induced/ MY-COMP inhibited genes, including ARHGAP11A, BUB1 and DEPDC1 which were not investigated by us as YAP-MMB target genes before and independently confirmed by RT-qPCR that MY-COMP effectively counteracts the YAP-mediated activation of these genes (Figure 3H).

**Figure 3:**
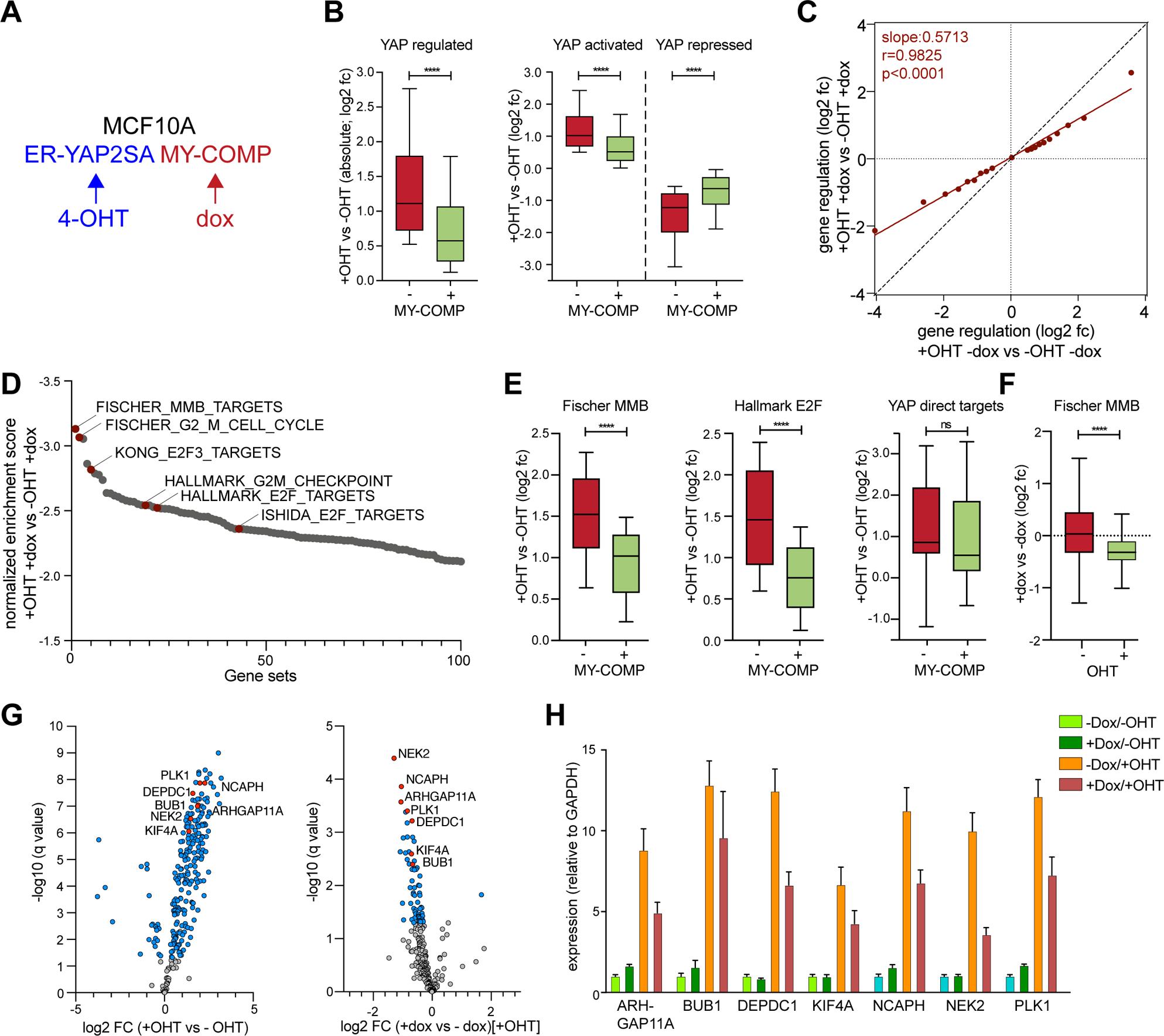
Global effects of MY-COMP on the YAP-dependent transcriptional program. A) Scheme for MCF10A cells stably expressing ER-YAP2SA and doxycycline-inducible MY-COMP B) ER-YAP2SA/MY-COMP cells were treated with and without 4-OHT and with and without doxycycline. RNA was isolated and subjected to RNA-seq. Box plots documenting global effect of MY-COMP on YAP-mediated gene activation and repression. The plot on the left shows absolute values for all significantly YAP-regulated genes (padj < 0.01), plots on the right show the effect of MY-COMP on YAP-dependent gene activation and repression. Box plots extends from the 25th to 75th percentiles. A horizontal line indicates the median, whiskers extend to 10-90% percentile and outliers are not shown. P-values were calculated using two sided, paired Wilcoxon signed-rank tests. Number of genes in each gene set: YAP regulated: 5,421. YAP activated: 2,830. YAP: repressed: 2,591. C) Correlation of YAP-dependent gene regulation in cell expressing and not expressing MY-COMP. Shown are the log_2_ fold-change (fc) values of gene regulation in cells after 4-OHT mediated activation of ER-YAP2SA. All YAP-regulated genes (padj<0.01) were sorted for YAP regulation and binned into 20 bins with 271 genes each. A linear model was used to calculate a regression line. r^2^, r squared correlation coefficient. D) Gene set enrichment analyses (GSEA) of RNA-seq data. Displayed are the normalized enrichment scores (NES) of the top 100 gene-sets downregulated by MY-COMP in the presence of 4-OHT. E) Box plots documenting the effect of MY-COMP on YAP-mediated activation of the following gene set. Fischer MMB targets: MMB target genes from https://doi.org/10.1093/nar/gkw523 (172 genes). Hallmark E2F targets: https://www.gsea-msigdb.org/gsea/msigdb/cards/HALLMARK_E2F_TARGETS (169 genes). YAP bound: Genes with YAP peak in +-1kb around TSS YAP binding, sorted for %FDR->sort decreasing and select 100 most significant (=strongest bound) from ^17^ (36 genes). Box plots were drawn as described in panel A. F) Box plots documenting the effect of MY-COMP on MMB-target genes without (-) and with (+) activation of ER-YAP2SA by 4-OHT. G) Volcano blots of RNA-seq data after induction of ER-YAP2SA (left) or after simultaneous expression of MY-COMP (right). q-values and fold change of MMB-target genes are plotted. 3 biological replicates per condition. H) Activation by ER-YAP2SA and repression by MY-COMP of the indicated genes selected out of the shown in E was verified by RT-qPCR.

### MY-COMP inhibits YAP-driven liver tumorigenesis

To assess the effect of MY-COMP on YAP-induced tumorigenesis *in vivo*, we used an autochthonous model of liver cancer. To trigger liver tumorigenesis, we performed hydrodynamic tail vein injection (HDTVI) to deliver transposon-based vectors expressing hyperactive YAP5SA mutant together with myristoylated AKT (myr-AKT) into livers of C57Bl6 mice. YAP5SA is a constitutive active YAP similar to the above described YAP2SA mutant but with three additional LATS1/2 sites mutated from S to A. The expression of YAP5SA and myr-AKT results in formation of multifocal hepatocellular carcinoma ^23^. To assess the effect of MY-COMP on YAP-induced tumorigenesis, we co-expressed myristolated AKT (myr-AKT) together with YAP5SA vectors expressing either GFP or MY-COMP (Figure 4A). Subsequently, we closely monitored the mice for tumor development. While myr-AKT and YAP5SA robustly induced liver tumors after 6 weeks, we found a significant reduction in hepatocellular carcinoma formation due to the presence of MY-COMP in this model (Figure 4B-D). MY-COMP expression decreases the liver to body weight ratio to levels that were observed in not injected animals of a similar age (Figure 4C). Quantification of the tumor area from H&E-stained liver sections revealed that the tumor burden of MY-COMP expressing livers is significantly reduced compared to GFP controls (Figure 4B,D). Taken together these data highlight the importance of the YAP-B-MYB interaction in YAP-driven tumor development.

**Figure 4:**
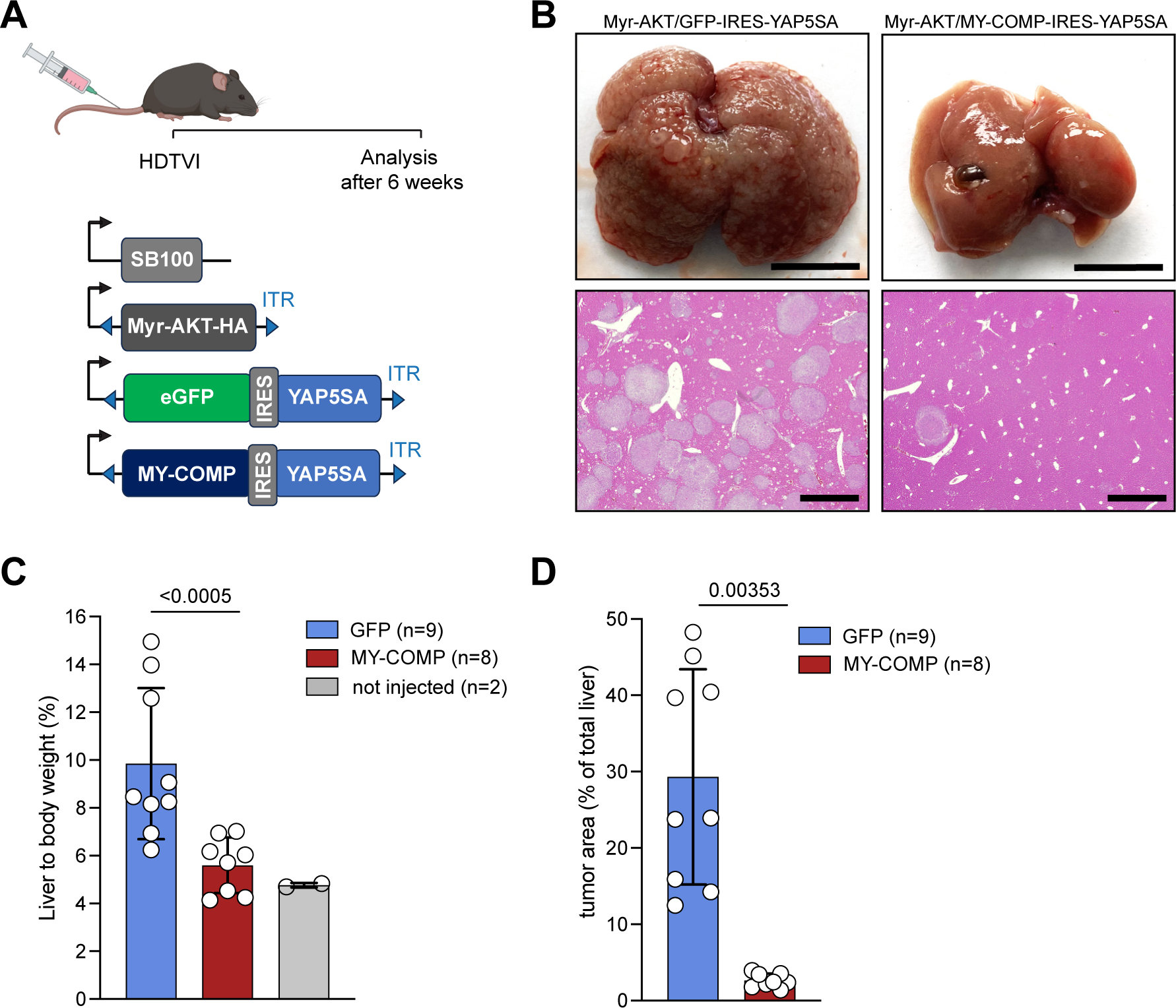
MY-COMP inhibits YAP-driven liver tumorigenesis. A) Scheme of the HDTVI-based model of liver tumorigenesis. Vectors encoding Myr-AKT and GFP-IRES-YAP5SA or Myr-AKT and MY-COMP-IRES-YAP5SA were injected along with the SB transposase into the tail vein of C57BL/6 mice. Liver tumor formation was assessed 6 weeks later. SB100=sleeping beauty transposase, ITR=inverted terminal repeat B) Top: Representative images of whole livers from mice injected with the indicated constructs. Bottom: Representative H&E images of the two groups. Scale bars: 1 cm C) Ratio of liver to body weight of the indicated groups. Welch t-test. D) Quantification of tumor area relative to the whole liver area. Welch t-test. GFP (n= 9), MY-COMP (n=8).

### MMB mediates oncogenic functions of YAP in uveal melanoma cell lines

To investigate whether MY-COMP can counteract the oncogenic functions of YAP in a cellular model for tumorigenesis, we next focused on YAP-dependent uveal melanoma (UM) cell lines. UM is the most frequent ocular malignancy in adults with a very poor prognosis. A key oncogenic event in uveal melanoma is the activation of YAP, resulting from mutations in GNAQ or GNA11, which encode α subunits of the Gq family of G proteins ^6^. This distinguishes uveal melanoma from cutaneous melanoma, which typically harbor mutations in B-RAF or N-RAS. The activation of YAP by GNAQ or GNA11 renders YAP a promising target for the treatment of uveal melanoma.

Immunostaining of a panel of different GNAQ/11 mutated uveal melanoma cell lines revealed prominent nuclear localization of YAP, while B-RAF mutated cutaneous melanoma cell lines exhibited predominantly cytoplasmic YAP staining (Figure 5A). Given the YAP-dependency of uveal melanoma, we investigated whether these cells would display increased sensitivity to MY-COMP-mediated YAP-B-MYB inhibition compared to B-RAF-mutated cell lines. To accomplish this, we introduced doxycycline-inducible MY-COMP into uveal melanoma and B-RAF-mutated melanoma cell lines, followed by the assessment of the optimal doxycycline concentration for robust MY-COMP expression. A concentration of 0.5 µg/µl was chosen for subsequent experiments (Supplemental Figure S2). After 72 hours of MY-COMP induction, flow cytometry analysis revealed a significant increase in the number of uveal melanoma cells with >4n DNA content, while no significant effect was observed in B-RAF-mutated cutaneous melanoma cells (Figure 5B,C). To further evaluate the role of the YAP-MMB interaction and the effect of MY-COMP on oncogenic transformation of uveal melanoma cells, we analyzed anchorage-independent growth in soft agar. Depletion of YAP/TAZ reduced the number of colonies in soft agar, as expected from previous studies (Figure 5D,E,F) ^6^. Similarly, RNAi mediated depletion of B-MYB or of LIN9, a core subunit of MuvB, strongly suppressed colony formation of 92.1 cells indicating the requirement of these MMB subunits for oncogenic transformation. Notably, expression of MY-COMP also significantly inhibited colony formation of 92.1 cells, suggesting that the YAP-B-MYB interaction is crucial for anchorage-independent growth of these cells (Figure 5D,E,F).

**Figure 5:**
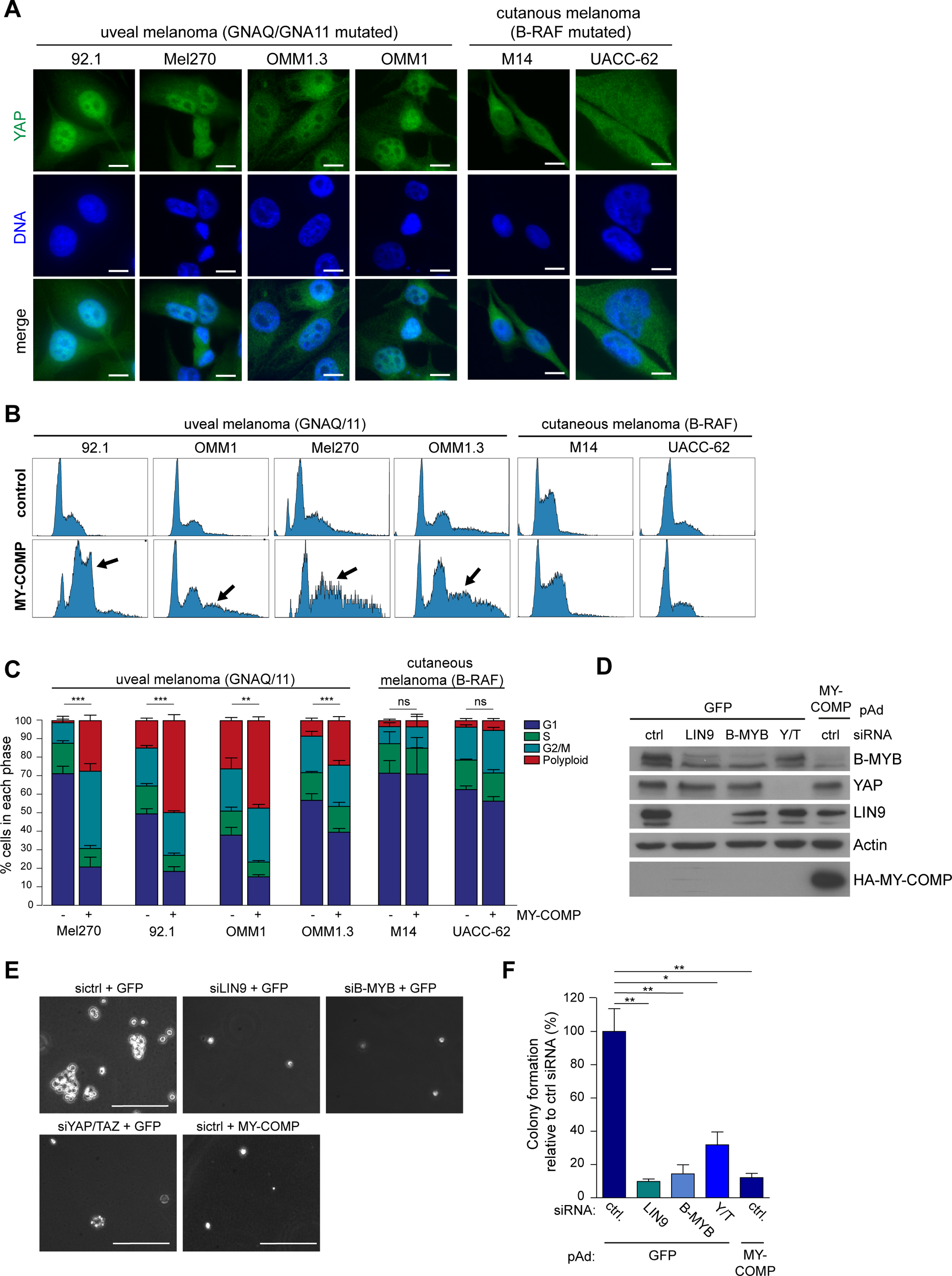
Uveal melanoma cells are sensitive to inhibition of the YAP-B-MYB interaction. A) A panel of uveal melanoma cell lines (92.1, Mel271, OMM1.2 and OM1) and B-RAF mutated cutaneous melanoma cell lines (M14 and UACC-62) were immunostained for YAP. Scale bar: 10 µm. YAP localizes to the nucleus in uveal melanoma cell lines but not in B-RAF mutated cutaneous cell lines. B) The cell lines described in A were stably transfected with doxycycline-inducible MY-COMP. MY-COMP was induced for 4 days and the fraction of cells in the different phases of the cell cycle was analyzed by FACS. C) Quantification of the FACS data obtained from three independent experiments. Error bars depict SEM. Student’s t-test was used to calculate the significance of the differences in polyploid cells. **=p<0.01, ***=p<0.001, ns= not significant. D) 92.1 cells were transfected with the indicated siRNAs directed at LIN9, B-MYB or YAP/TAZ or were infected with a recombinant adenovirus encoding MY-COMP. As control, cells were transfected with a siRNA against luciferase and infected with an adenovirus encoding GFP. Expression of the indicated proteins was investigated by immunoblotting. Actin served as a control. E) and F) To assay for anchorage-independent growth, cells as described in D were seeded in 0.6 % agarose and grown for 21 days. E shows representative images (scale bar: 25 µm) and F shows a quantification of one experiment performed in triplicate. Error bars indicate SEM.

We next investigated the effect of MY-COMP on the expression of MMB target genes in 92.1 cells by performing RNA-seq. The volcano plot in Figure 6A demonstrates that expression of MY-COMP resulted in changes in MMB-target gene expression, with the majority of MMB target genes downregulated and a smaller fraction upregulated. Similarly, siRNA mediated depletion of YAP and TAZ also downregulated MMB target genes (Figure 6B). Notably there exists a strong correlation between significantly downregulated genes by MY-COMP and by YAP/TAZ knockdown (Figure 6C). Gene Set Enrichment Analyses (GSEA) confirmed that both MY-COMP and the depletion of YAP/TAZ strongly inhibit the expression of MMB-target genes with a normalized enrichment score of -2.16 and -3.21, respectively (Supplemental Figure S3). To identify direct targets of YAP in uveal melanoma cells, we performed CUT&RUN experiments and identified 7,457 high-confidence peaks (Figure 6D). Notably, YAP bound mainly to distal intergenic and intronic regions, and to a lesser extend to promoters (10% of peaks). Because a large fraction of YAP binding sites were gene distal, we ask whether they correspond to enhancer regions. To identify enhancers, we performed CUT&RUN for histone H3 mono- and trimethylated at lysin 4 (H3K4me1 and H3K4m3). Our analysis revealed that out of a total of 13,211 enhancers regions defined by H3K4me1-positive/ H3K4me3-negative regions that are not within 1 kb of a transcription start site, 1,912 (14.5%) displayed one or more YAP peaks (Figure 6E). This indicates that YAP binds to a significant proportion of all enhancers in 92.1 cells. Moreover, CUT&RUN for acetylated histone H4 and for histone H3 acetylated at lysin 27 (H3K27Ac) showed that YAP-bound enhancers exhibit high levels of acetylation and are thus active (Figure 5F). Notably, the distribution of H4Ac and H3K27Ac at YAP-peaks is not random; instead, we observed the highest enrichment of acetylated histones precisely at the center of the YAP peak. By integrating the RNA-seq and CUT&RUN data, we found that out of the 3,020 downregulated genes observed upon YAP/TAZ depletion, 1,468 were direct YAP-regulated genes, as indicated by an associated YAP-peak (Figure 6G). Intriguingly, 381 of these direct YAP-target genes were also found to be downregulated following the expression of MY-COMP (Figure 6H). Further examination through GO analysis of these 381 genes showed enrichment in categories associated with essential cellular processes such as cell division, cell cycle regulation, chromosome segregation, Hippo signaling, transcription, and DNA-repair (Figure 6I). Examples for YAP/ MMB regulated genes from this group are NEDD9 and CDC25B. Genome browser tracks illustrating CUT&RUN data of YAP binding to active enhancers of these genes are shown in Figure 6J.

**Figure 6:**
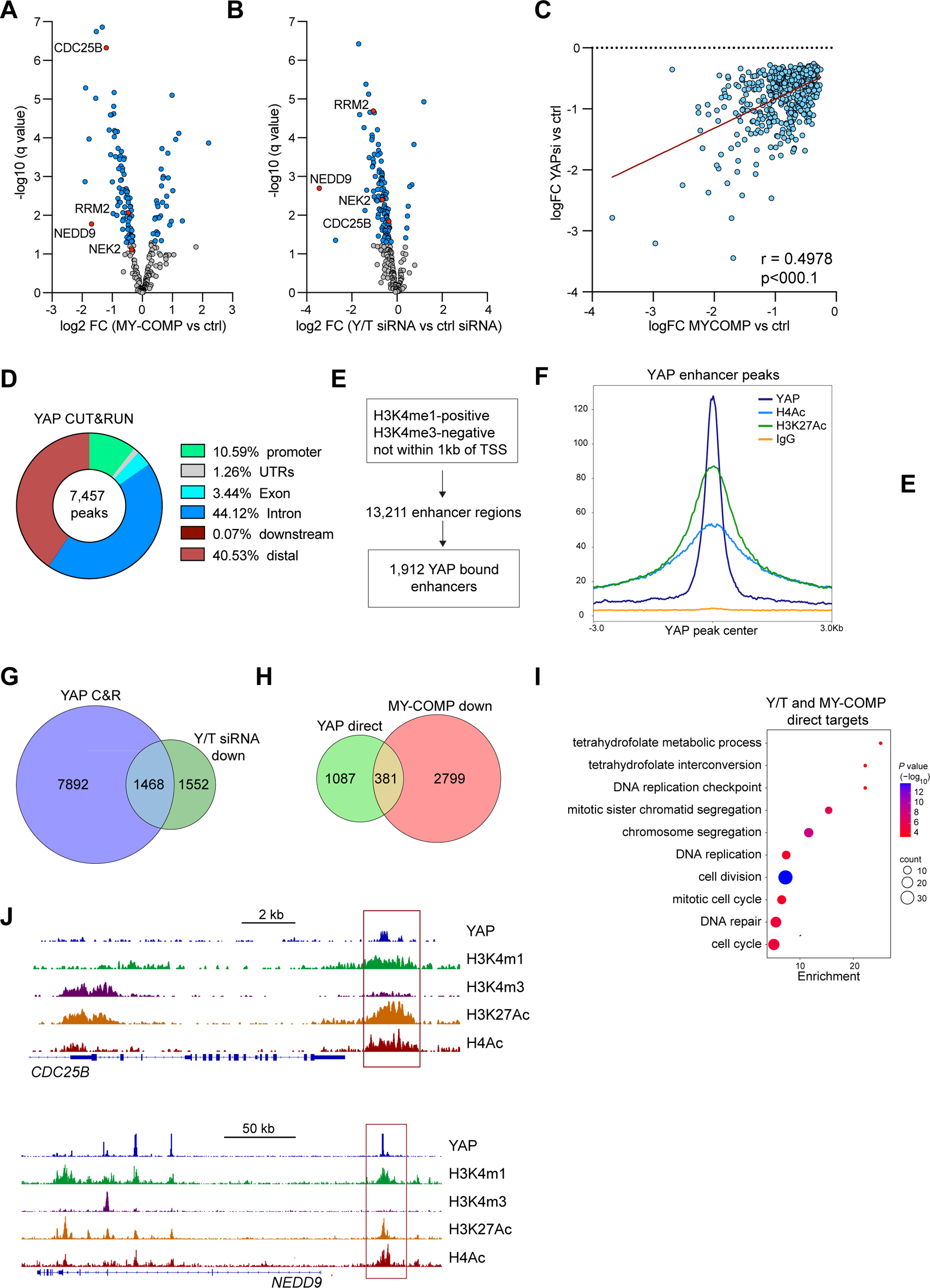
YAP regulates the expression of MMB target genes in uveal melanoma cells by binding to distant enhancers. A) Volcano blots of RNA-seq data after expression of MY-COMP-cells in 92.1 cells. q-values and fold changes of MMB-target genes are plotted. 3 biological replicates per condition. B) Volcano blots of RNA-seq data after siRNA mediated depletion of YAP/TAZ. q-values and fold changes of MMB-target genes are plotted. 3 biological replicates per condition. C) Correlation of downregulated genes by MY-COMP with genes downregulated by YAP/TAZ depletion. D) CUT&RUN was performed to determine the genome wide distribution of YAP in 92.1 cells. The distribution of CUT&RUN peaks for YAP relative to known genes in the genome is shown. E) Identification of 13,211 enhancers in 92.1 cells by CUT&RUN for H3K4me1 and H3K4me3. By comparison with YAP CUT&RUN, 1,912 YAP bound enhancers were identified. F) Line plot of enrichment of IgG, YAP, H3K27Ac and H4Ac at YAP enhancer binding sites. G) Integration of YAP CUT&RUN data and RNA-seq data. Of 3,020 genes significantly downregulated after the depletion of YAP/TAZ, 1,468 are direct YAP-target genes with a nearby YAP CUT&RUN peak. H) Venn diagram showing that of the 1,468 direct YAP-targets, 381 are also downregulated by MY-COMP. I) GO analysis of 381 YAP-target genes downregulated by MY-COMP. J) Genome browser CUT&RUN tracks of CDC25B and NEDD9 (K) demonstrating binding of YAP to a nearby, active enhancer (red box).

### NEK2 is a clinically relevant YAP/ MMB target in uveal melanoma

In light of our finding that transformation of uveal melanoma cell lines depends on YAP and can be inhibited by MY-COMP, we aimed to identify YAP-MMB-target genes that may have clinical relevance for uveal melanoma. To address this, we asked whether the expression of YAP-MMB-target genes identified in the RNA-seq and CUT&RUN analysis correlated with the overall survival of uveal melanoma patients. We stratified patients into groups based on the expression quintiles of the corresponding gene in the TCGA data set. Through this analysis we discovered that Never in Mitosis (NIMA) Related Kinase 2 (NEK2), one of the above identified targets of YAP-MMB in uveal melanoma cells, exhibited a significant association with poor survival in uveal melanoma patients but not in cutaneous melanoma patients (Figure 7A, B, Supplemental Table 3). This finding indicates that NEK2 is one of the YAP-MMB-regulated genes with clinical relevance specifically for uveal melanoma, highlighting its potential as a therapeutic target or prognostic marker.

**Figure 7:**
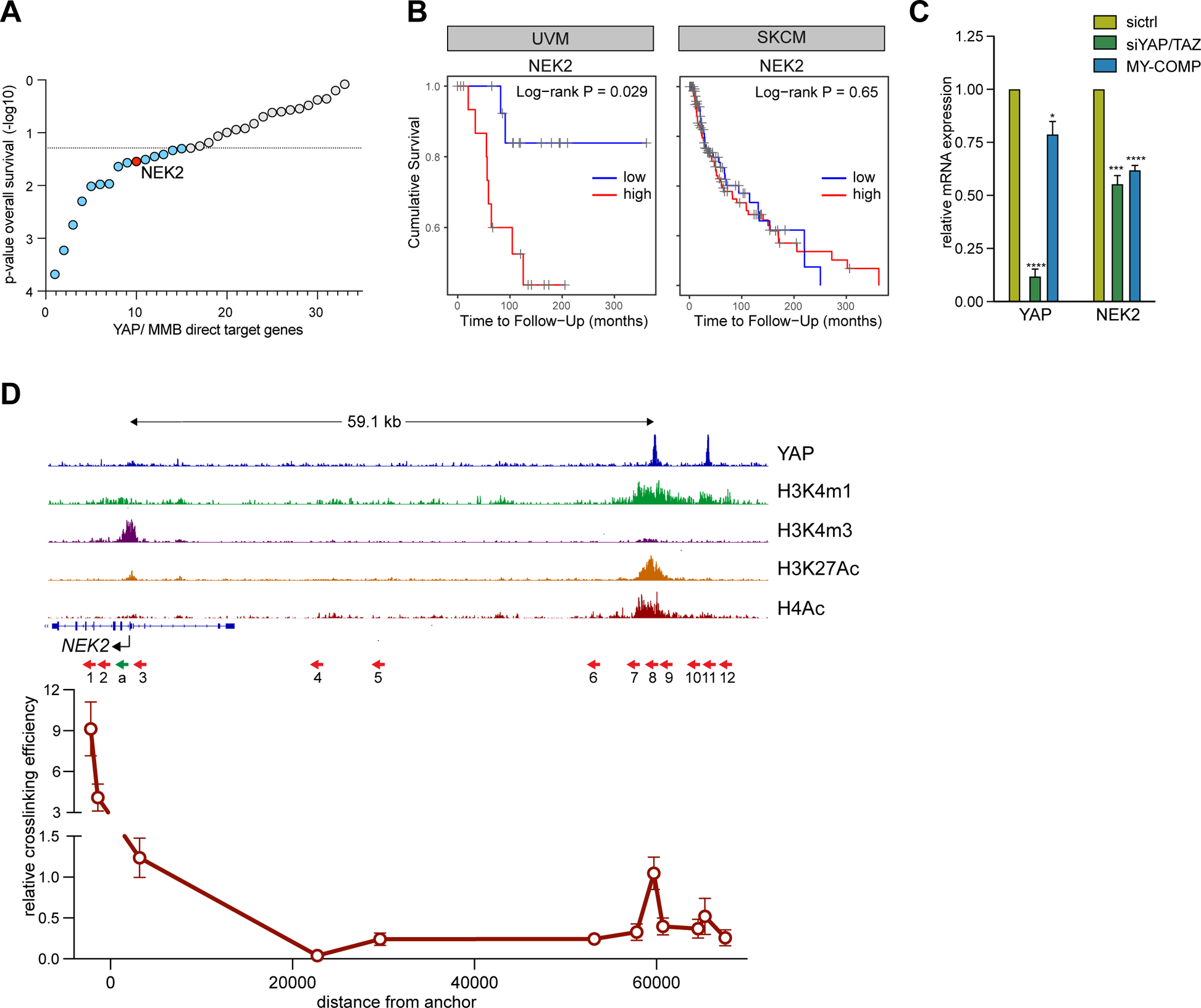
NEK2 is a clinically relevant YAP/ MMB target in uveal melanoma. A) The significance for association with overall survival (OS) of uveal melanoma patients was calculated by stratifying for expression quintiles of YAP/MMB target genes. This analysis identified NEK2 as an MMB-target gene whose high expression is significantly associated with poor survival of uveal melanoma patients. B) Kaplan-Meier survival analysis of uveal melanoma (UVM) and skin cutaneous melanoma patients (SKCM) in the TCGA datasets stratified by quintile NEK2 expression. C) The expression of YAP and NEK2 after siRNA-mediated depletion of YAP/TAZ or expression of MY-COMP in 92.1 cells was analyzed by RT-qPCR. N=3 replicates. Error bars indicate SD. Student’s t-test. D) 3C-qPCR chromatin capture experiments were performed to assess the interaction between the YAP bound enhancer and the NEK2 promoter. The top shows the CUT&RUN tracks of YAP, H3K4me1, K3K4me3, H4Ac and H3K27Ac at the NEK2 locus demonstrating binding of YAP to an active enhancer 60 kb telomeric of the NEK2 promoter. The arrows indicate the location of primers used in the 3C-qPCR analysis. The constant anchor primer (a) is indicated in green. Interaction frequencies were determined by qPCR after HindIII digestion and re-ligation relative to the ligation products from the BAC DNA covering the same genomic region. Data shown are from one representative experiment from two independent experiments.

We confirmed by RT-qPCR MY-COMP as well as the depletion of YAP/TAZ leads to the downregulation of NEK2 expression (Figure 7C). To understand how YAP regulates NEK2, we analyzed our CUT&RUN data and found that YAP binds to an active enhancer approximately 60kb telomeric of *NEK2* (Figure 7D). To confirm the long-range interaction between the NEK2 promoter and the YAP-bound enhancer, we performed chromatin conformation capture (3C) experiments. We used a fixed anchor primer in the NEK2 promoter and a set of 11 specific primers spanning the whole genomic region of more than 60 kb between the NEK2 promoter and the YAP-bound enhancer (Figure 7D). Regions close to the anchor exhibited high interaction frequencies, as expected. The interaction frequency was reduced in the genomic region between the promoter and the enhancer. Importantly, interaction frequencies strongly increased again at the YAP-bound NEK2 enhancer. This confirms the direct interaction between the NEK2 promoter and the YAP-bound enhancer by chromatin-looping, providing additional evidence for the regulatory relationship between YAP and NEK2 and suggesting that YAP binding to the enhancer region contributes to NEK2 gene expression.

### Uveal melanoma cells are dependent on NEK2 for transformation and survival

Next, to test whether NEK2 is a downstream target contributing to the oncogenic properties of YAP in UM cells, we employed the NEK2 inhibitor INH1, which is known to block the interaction of NEK2 with HEC1 triggering the degradation of NEK2 ^24^. Interestingly, UM cell lines were more sensitive towards NEK2 inhibition compared to CM cells (Figure 8A). Similarly, while high doses of INH1 (25µM) affected proliferation of both UM and CM cell lines, inhibition of NEK2 with a lower concentration of INH1 (12.5µM) more strongly affected proliferation of UM cell lines, whereas growth of CM was not significantly inhibited by this concentration of INH1 (Figure 8B,C). To independently validate the effects of NEK2 loss, we generated individual UM cell lines stably expressing one of two different doxycycline-inducible NEK2-specific shRNAs (Supplemental Figure S4A). Doxycycline-mediated depletion of NEK2 by the two shRNAs was validated by RT-qPCR and immunoblotting (Supplemental Figure S4B,C). Consistent with the experiments with INH1, doxycycline-induced silencing of NEK2 resulted in a dose dependent growth-deficit of UM cell lines (Supplemental Figure S4D). Furthermore, depletion of NEK2 reduced anchorage-independent growth of colonies in soft agar (Figure 8D). Inhibition of NEK2 culminated in apoptotic cell death, indicating that NEK2 is required for survival of uveal melanoma cells (Figure 8E and Supplemental Figure S5).

**Figure 8:**
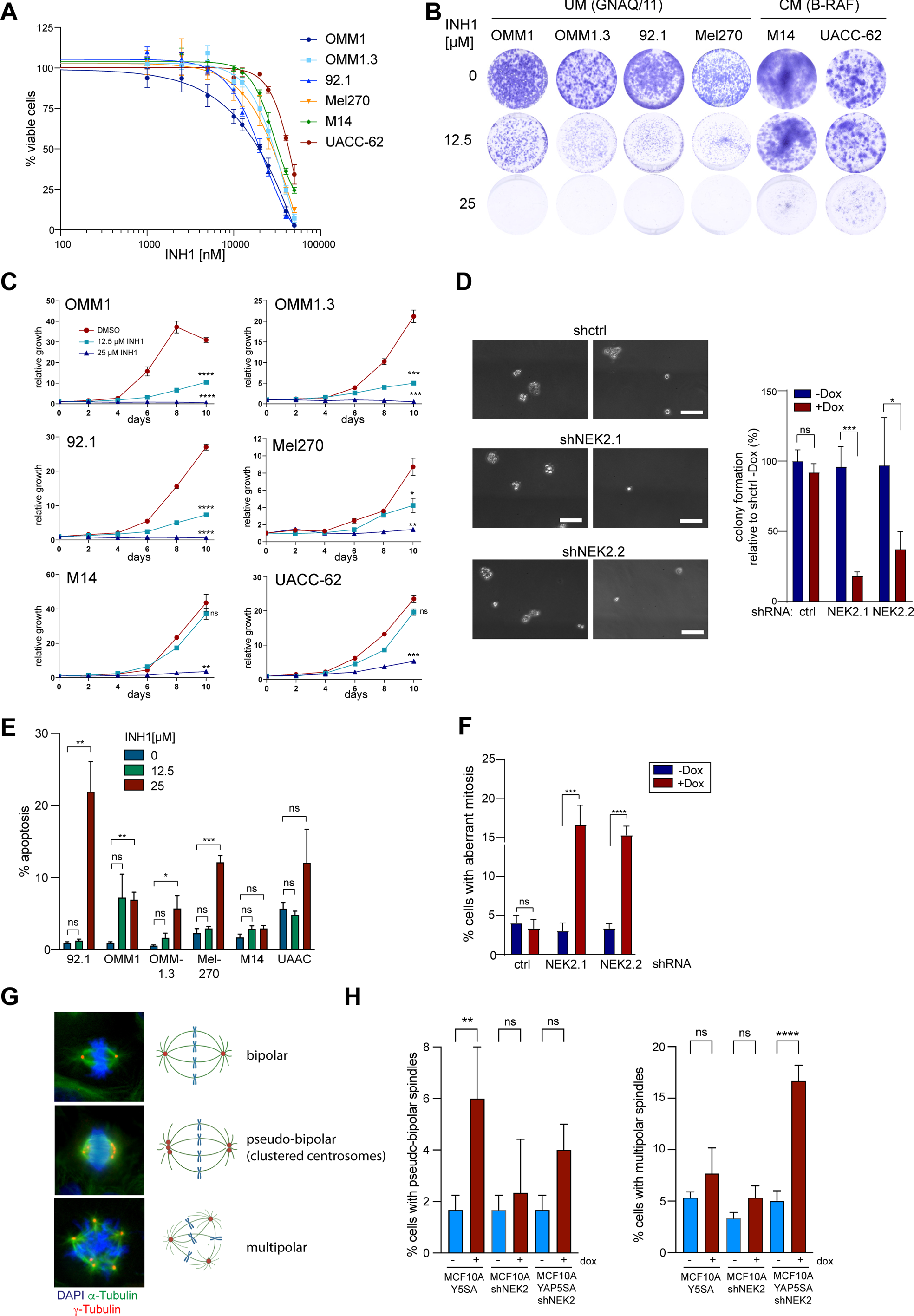
Uveal melanoma cells depend on NEK2 for transformation and survival. A) UM and CM cells were treated with the indicated concentrations of INH1 and cell viability was analyzed by MTT assay. IC-50 values for the individual cell lines were calculated using a nonlinear regression (curve fit). B) and C) UM and CM cells were seeded at low density and left treated or untreated with the indicated concentrations of INH1 for 10 days. Cells were stained with crystal violet. Representative images are shown in B. Quantification of crystal violet staining is shown in C. n=3 biological replicates each performed in triplicates. Error bars indicate SEM of one representative experiment. Student’s t-test. *=p<0.05, **=p<0.01, ***=p<0.001, ****=p<0.0001, ns: not significant (refers to cells at day 10). D) 92.1 cells transfected with the indicated shRNAs were seeded in 0.3 % agarose and grown with and without doxycycline for 21 days to measure anchorage independent cell growth. Representative images (scale bar: 100 µm) quantification of one experiment performed in triplicates is shown. Error bars indicate SD. Student’s t-test. *=p<0.05, ***=p<0.001, ns: not significant. E) UM and CM cells were treated for 4 days with the indicated INH1 concentrations. Cells in subG1 phase were quantified by FACS. n=3 biological replicates. Error bars indicate SEM. Student’s t-test. *=p<0.05, **=p<0.01, ***=p<0.001, ns: not significant. F) 92.1 with the indicated shRNAs were treated with and without doxycycline. Mitotic spindles were investigated by staining for DAPI and α-tubulin and cells with aberrant mitosis were quantified. N=3. G) MFC10A cells expressing doxycycline-inducible YAP5SA, doxycycline-inducible NEK2 shRNA, or expressing both doxycycline-inducible YAP5SA and NEK2 shRNA were treated with 72 hours with 1 µg/ml doxycycline or with solvent. Mitotic spindles were analyzed by immunostaining for α-tubulin (green), γ-tubulin(red) and Hoechst (blue). Representative immunofluorescence images are shown. H) Frequencies of cells with pseudo-bipolar and multipolar spindles before and after induction of YAP5SA and/ or the NEK2 specific shRNA for 72 hours. n=3 independent experiments. Significance was determined by one-way ANOVA with Sidak multiple comparison; ** p<0.01; ****p ≤ 0.0001.

### NEK2 suppresses multipolar spindle formation in YAP-dependent tumor cells

NEK2 is a mitotic kinase that is involved in centrosome duplication and separation, assembly of the mitotic spindle and in the spindle assembly checkpoint ^25^. Depletion of NEK2 in 92.1 uveal melanoma cells resulted in significant increase in multipolar mitotic spindles, in contrast to control cells in which most of the spindles were bipolar (Figure 8F). To explore the impact of NEK2 depletion in relationship to YAP, we shifted our focus to untransformed MCF10A cells and established cell lines expressing doxycycline-inducible YAP5SA, doxycycline-inducible NEK2 shRNA, or both. Upon analyzing the mitotic spindles by staining for α-tubulin and γ-tubulin, we observed that the expression of YAP5SA led to an increased proportion of cells with clustered pseudo-bipolar spindles and a moderate increase in cells with multipolar spindles (Figure 8G, H). Depleting NEK2 in MCF10A cells that do not overexpress YAP5SA did not change the percentage of cells with multipolar spindles or clustered centrosomes. Strikingly, however, when NEK2 was depleted in cells expressing YAP5SA, this led to a substantial increase in the formation of multipolar spindles coupled to a decrease in centrosome clustering (Figure 8H). These data indicate that NEK2 suppresses the formation of multipolar mitosis by oncogenic YAP by promoting centrosome clustering. In summary, our experiments identify NEK2 as a downstream target of YAP-MMB that is required for proliferation, transformation and survival of YAP-dependent tumor cells.

## DISCUSSION

Aberrant activation of YAP is frequently observed in human cancer and is generally correlated with a poor outcome ^26^. Targeting YAP and TAZ represents a potential therapeutic opportunity. Since YAP and TAZ exert their effects on gene expression mainly through their interaction with TEAD transcription factors, disrupting the TEAD-YAP interaction is an obvious approach inhibiting YAP/TAZ. Various studies have reported the efficacy of peptide-based antagonists in inhibiting YAP activity by interfering with the YAP-TEAD interaction ^27^. In addition, several drugs that target the palmitoylation of TEAD have been developed and demonstrated anti-tumorigenic properties in preclinical studies ^28, 29,30^. Overall, targeting the YAP-TEAD interaction through peptide-based antagonists or drugs that inhibit TEAD palmitoylation represent promising strategies for cancer therapy. A potential drawback of inhibiting TEAD is that TEAD has different coactivators and corepressors that modulate its activity. One of the corepressors is VGLL4 (Vestigial-like family member 4), which can repress TEAD-mediated transcription ^31–33^. When TEAD is inhibited, it not only affects the activation of certain TEAD target genes but also leads to the repression of TEAD-VGLL4 complexes. Inhibiting TEAD may therefore result in decreased activity of VGLL4, which can lead to the loss of its repressive effects on TEAD targets. Consequently, this loss of repression may inadvertently activate certain TEAD target genes.

Here we used an approach to interfere with the binding of B-MYB to YAP. YAP interacts with B-MYB, a subunit of the MMB complex, to co-regulate a set of cell cycle genes ^17, 18^. Our previous biochemical characterization of the YAP-B-MYB interaction demonstrated that a central domain of B-MYB directly interacts with the WW-domains of YAP. Building on this observation we have created a YAP-binding construct named MY-COMP, which contains the YAP-binding domain of B-MYB fused to a nuclear localization signal. We have previously shown that MY-COMP prevents the YAP-induced mitotic entry of murine cardiomyocytes by blocking the binding of B-MYB to YAP ^18^.

We now demonstrate that MY-COMP is also effective in human cancer cells, where it inhibits the interaction of B-MYB and YAP, thereby interfering with mitosis and cytokinesis and preventing transformation. MY-COMP reverted the gene expression program induced by oncogenic YAP in MCF10A cells, with E2F- and MMB-target genes being particularly responsive to MY-COMP, consistent with the intended design. Notably, MY-COMP had significantly weaker effects on endogenous MMB-target gene expression in normal pre-malignant MCF10A cells, suggesting that blocking the WW-domains by MY-COMP could result in a relative low toxicity.

The WW-domain of YAP serves as a binding module that interacts with specific PPXY motifs in proteins that either promote cell proliferation or suppress YAP activity ^8, 34^. For example, the WW-domain mediates interaction with inhibitors of YAP activity, like AMOTL1 or LATS1/2 that sequester YAP in the cytoplasm and prevent translocation into the nucleus. Most of the WW-domain mediated interactions that block YAP activity are cytoplasmic, whereas transcription factors, such as B-MYB, exclusively interact with the YAP WW-domain in the nucleus. MY-COMP was designed to suppress the nuclear activity of YAP, which was achieved by fusion to a nuclear localization signal.

As a downstream target of YAP by which MY-COMP might prevent transformation of YAP-dependent tumors, we identified NIMA-related kinase (NEK2), a mitotic serine/threonine kinase that localizes to the centrosomes and to the kinetochore, as a candidate. NEK2 plays a critical role in centrosome separation, and bipolar spindle formation ^35,36^. While NEK2 is essential for cell cycle progression of normal cells, it is frequently overexpressed in cancer, thereby potentially opening a specific therapeutic window. Our data indicate that overexpression of oncogenic YAP is associated with the formation of multipolar mitosis and that NEK2 acts as a suppressor of this phenotype. Cancer cells cluster their excess centrosomes, resulting in the formation of pseudo-bipolar spindles that effectively separate chromosomes ^37^. Disrupting centrosome clustering leads to the creation of multipolar spindles, which are linked to the mis-segregation of chromosomes and ultimately result in cell death ^37^. NEK2 is primarily recognized for its role in inactivating centrosome cohesion during the G2 phase by phosphorylating linker proteins, resulting in the disassembly of the centrosome linker. However, a recent study has reported the suppression of the multipolar mitosis phenotype in p53-negative breast cancer cells by NEK2, which is consistent with the findings presented here^38^. These observations underscore the potential of NEK2 as a therapeutic target for controlling abnormal mitosis associated with YAP overexpression. In this regard, multiple preclinical studies have demonstrated that targeting NEK2 can inhibit tumor growth ^24, 39, 40^. In conclusion, strategies aimed at suppressing YAP-regulated genes through nuclear blockade of the WW-domains, as demonstrated by MY-COMP, or targeting specific YAP-MMB regulated genes such as NEK2, hold promise as novel approaches to counteract the pro-tumorigenic functions of YAP.

## MATERIAL AND METHODS

### Cell lines

MCF10A cells were cultured in DMEM/F-12 supplemented with 5% horse serum, 1% penicillin/ streptomycin, 10 µg/ml insulin, 500 ng/ml hydrocortisone, 20 ng/ml EGF and 100 ng/ml cholera toxin. Uveal melanoma cell lines were maintained in RPMI and cutaneous melanoma cells and HeLa cells were cultured in DMEM supplemented with 10% FCS (Thermo Fisher Scientific) and 1% penicillin/streptomycin (Thermo Fisher Scientific).

### Hydrodynamic tail vein injection (HDTVI)

For intrahepatic delivery of the sleeping beauty transposon system, 6-week-old male mice (Janvier) were anesthetized with isoflurane and hydrodynamically injected with 10µg SB100X, 25µg pSBbi-w/o-puro-myr-AKT-HA, 25µg of either pSBbi-w/o-puro-GFP-IRES-YAP5SA or pSBbi-w/o-puro-MY-COMP-IRES-YAP5SA. Animal experiments were approved by the state of Thuringia under the animal license FLI-21-015. The mice were kept in individually ventilated cages (IVCs) under Specific Pathogen Free (SPF) conditions with a 12 h/12 h dark/light cycle at a temperature of 20 °C and a relative humidity of 55% according to the directives of the 2010/63/EU and GV SOLAS. All plasmids were diluted in Ringer-Lactat solution and adjusted to 10% of the total body weight.

### Adenovirus

Adenoviral constructs were described previously ^18^, using the ViralPower adenoviral expression system (Thermo Fischer Scientific). Adenoviruses were titered using the Adeno-X Rapid Titer Kit (TaKaRa).

### Plasmids

For HDTVI the following plasmids were used: pCMV(CAT)T7-SB100 containing SB100X transposase (Addgene #34879). For Myr-AKT and YAP5SA expression the pSBbi-Puro vector from Addgene (#60523) was used. The Puromycin cassette was removed from the vector by digestion with EcoRI and BsrGI. Myristolated AKT was amplified by PCR from pT3-myr-AKT-HA (Addgene #31789) and cloned into the pSBbi-w/o-Puro backbone. For the GFP-IRES-YAP5SA construct, GFP was amplified by PCR from pLeGO-iG2 (Addgene #27341). The IRES site was amplified by PCR from pInducer-21 (Addgene #46948). YAP5SA was amplified by PCR from Addgene #33093. Gibson assembly was used to clone the amplified genes into pSBbi-w/o-Puro expression vector. For the MY-COMP-IRES-YAP5SA MY-COMP was amplified by PCR from pCDNA4/TO-MY-COMP vector ^18^. HDTVI plasmids were purified using the endotoxin free NucleoBond PC 500 EF kit.

### Lentiviral production and infection

Lentiviral particles were generated in HEK293T co-transfected with psPAX2, pCMV-VSV-G and a lentiviral vector. Filtered viral supernatant was diluted 1:1 with culture medium and supplemented with 4 μg/ml polybrene (Sigma). Infected cells were selected 48 h after infection with the appropriate antibiotics for 7 days.

### Proliferation assays

Cells were plated in 24-well plates. At the indicated time points, cells were fixed in 10 % formalin and stained with 0.1 % crystal violet. The dye was extracted with 10% acetic acid and the optical density was determined as described ^41^.

### MTT assays

Cells were seeded in 96-well plate and treated the following day with the indicated concentrations of INH1 in complete medium. 20 µl of thiazolyl blue tetrazolium bromide (5mg/ml) was added directly to the medium and cells. After 2 h at 37°C, 100 μl DMSO was added, the plate was incubated for 20 minutes at RT and the absorbance was measured in a microplate reader at 590nm. The absorbance was normalized to the negative control of medium without cells, and the cell viability was calculated relative to DMSO control.

### RT-qPCR

Total RNA was isolated using peqGOLD TriFast (Peqlab). RNA was transcribed using 100 units RevertAid reverse transcriptase (Thermo Fisher Scientific). Quantitative real–time PCR reagents were from Thermo Fisher Scientific and real-time PCR was performed using a qTower3G (Analytik Jena). Expression differences were calculated as described before ^42^. Primer sequences are listed in Supplemental Table 1.

### Immunoblotting and immunoprecipitation

Cells were lysed in TNN (50mM Tris (pH 7.5), 120mM NaCl, 5mM EDTA, 0.5% NP40, 10mM Na4P_2_O_7_, 2mM Na_3_VO_4_, 100mM NaF, 1mM PMSF, 1mM DTT, 10mM β-glycerophosphate, protease inhibitor cocktail (Sigma)). Proteins were separated by SDS-PAGE, transferred to a PVDF-membrane and detected by immunoblotting with the first and secondary antibodies. For immunoprecipitation of FLAG-tagged proteins, protein G dynabeads (Thermo Fisher Scientific) were first coupled with 1 µg FLAG-antibody (Sigma, F3165) and then immunoprecipitated with 1mg whole cell lysate. After five times of washing with TNN, proteins were separated by SDS-PAGE and detected by immunoblotting using the desired antibodies. Antibodies are listed in Supplemental Table 2.

### Immunostaining

Cells were fixed with 3% paraformaldehyde and 2% sucrose in PBS for 10 minutes at room temperature. Cells were permeabilized using 0.2% Triton X-100 (Sigma) in PBS for 5 minutes and blocked with 3% BSA in PBS-T (0.1% Triton X-100 in PBS) for 30 minutes. Primary antibodies were diluted in 3% BSA in PBS-T and incubated with the cells for 1 hour at room temperature or overnight for cell cycle profile. After three washing steps with PBS-T, secondary antibodies conjugated to Alexa 488 and 594 (Thermo Fisher Scientific) and Hoechst 33258 (Sigma) were diluted 1:500 in 3% BSA in PBS-T and incubated with the coverslips for 30-60 minutes at room temperature. Finally, slides were washed three times with PBS-T and mounted with Immu-Mount™ (Thermo Fisher Scientific). Pictures were taken with an inverted Leica DMI 6000B microscope equipped with a Prior Lumen 200 fluorescence light source and a Leica DFC350 FX digital camera.

### Histology and Immunohistochemistry

Murine livers were fixed with 10% formalin (3.7% formaldehyde in 1 x PBS) for at least 48 hours at 4°C. Livers were then dehydrated and embedded in paraffin. Paraffin section (5µm) were stained with hematoxylin and eosin (H&E). Slides were imaged using the Axio Scan.Z1 microscope (Zeiss).

### siRNA transfection

Double-stranded RNA was purchased from Eurofins or Thermo Fischer Scientific. siRNAs were transfected using RNAiMAX (Thermo Fisher Scientific). siRNAs are listed in Supplemental Table 1.

### Flow cytometry

Samples were washed with ice cold PBS and fixed in 80% ice cold ethanol. Then, cells were washed with ice cold PBS and resuspended with 38 mM sodium citrate with 500 µg/ml RNase A for 30 min at 37°C. Before cells were analyzed on a Beckman Coulter Fc500, 43 mM propidium iodide was added.

### Soft agar assay

Cells were resuspended in 0.35% low melting agarose in soft agar medium [DMEM, 3.7% 1M sodium bicarbonate, 20% FCS, 20 mM glutamax, 9 mg/ ml D-glucose, 1% PenStrep]. 10,000 cells were seeded on a base layer of 0.7% low melting agarose in soft agar medium in a 6-well plate. For induction of the shRNA, 0.5 μg/ml doxycycline was added directly to the soft agar medium. Cells were cultivated for 14-or 18 days and fed every 3 days. Colony formation was analyzed by microscopy.

### RNA-Seq

Total RNA was isolated from three biological replicates. DNA libraries were generated using 1µg RNA with the magnetic mRNA isolation module and NEBNext Ultra II RNA Library Prep Kit for Illumina (New England Biolabs). DNA library was amplified by PCR and quality was analyzed using the fragment analyzer (Advanced Analytical). Libraries were sequenced on the NextSeq 500 platform (Illumina).

### CUT&RUN

CUT&RUN was carried out as described ^43, 44^. For CUT&RUN with H3K4m1, H3K4m3, H3K27Ac and H4Ac antibodies, a mild crosslinking with 1% formaldehyde was performed for 1min. Subsequently, digitonin was left out of all buffers for crosslinked samples. CUT&RUN for YAP was performed without crosslinking. Libraries were made with 6 ng of CUT&RUN DNA fragments using the NEBNext Ultra II DNA Library Prep Kit for Illumina (NEB #E7645S). The manufacturer’s protocol was adjusted to account for shorter DNA fragments as described previously ^45^.

### Quantitative Chromosome Conformation Capture Assay (3C-qPCR)

3C-qPCR was performed as described previously ^46^. Briefly, 92.1 cells were trypsinized, washed with PBS and resuspended in 1.5 ml Nucleus Buffer 1 (0.3M sucrose, 15 mM Tris-HCl pH 7.5, 6 0mM KCl, 15 mM NaCl, 5 mM MgCl_2_, 0.1mM EGTA, 0.5mM DTT, 0.1mM PMSF, protease inhibitor cocktail 1:1000). 0.5 ml Nucleus Buffer 2 (0.8% (v/v) IGEPAL CA 630 in Nucleus Buffer 1) was added. After incubation on ice for 3 min, 4 ml Nucleus Buffer 3 (1.2 M sucrose, 15 mM Tris-HCl pH 7.5, 60 mM KCl, 15mM NaCl, 5 mM MgCl_2_, 0.1mM EGTA, 0.5mM DTT, 0.1mM PMSF, protease inhibitor cocktail 1:1000) was added. Nuclei were crosslinked with 1% formaldehyde in 3C-buffer (50mM Tris-HCl pH 8, 50mM NaCl, 10mM MgCl_2_, 1mM DTT) for 10 min at room temperature, incubated with 0.125 M glycine for 7 min and washed with 1 ml 3C-buffer. Crosslinked nuclei were first resuspended in 100 µl 1x r2.1 restriction buffer (New England BioLabs) and then treated with 2% SDS for 1h at 37°C and with 1% Triton X-100 for 1h at 37°C. Samples were digested with 450 units of HindIII (ThermoFisher Scientific) overnight at 37°C. Samples were incubated with 1% SDS at 37°C for 1h, 3.67 ml 1x Ligation Buffer (50mM Tris-HCl pH 7.5, 10mM MgCl_2_, 1mM ATP, 10mM DTT). Afterwards, 1% Triton X-100 was added and the samples were incubated at 37°C for 2 h. Ligation was performed with 400 units of T4 DNA Ligase (New England Biolabs) at 16°C overnight. Samples were incubated with 1µg RNAse A (Sigma) for 1h at 37°C. Next, 2 ml PK-buffer (10mM Tris-HCl pH 8, 5mM EDTA, 0.5% SDS), 1.5ml ddH_2_O, and 100µg proteinase K (Sigma) were added and digested at 50°C for 1h. De-crosslinking was performed at 65°C for 4h. The 3C-library was purified using phenol/chloroform extraction and ethanol precipitation. Ligation products were quantified in qPCR reactions). To calculate primer efficiencies, a serial dilution of BAC clone RP11-122M14 (BACPAC Resources Center at BACPAC Genomics, Inc., Emeryville) spanning the NEK2 locus, digested with 450U HindIII and ligated with 2000U T4 DNA Ligase, was used. Values were normalized toward a loading control of a primer set not amplifying across a restriction site. Primer sequences are listed in Supplemental Table 1.

### Statistical Analysis

Base calling was performed with Illumina’s CASAVA software or FASTQ Generation software v1.0.0 and overall sequencing quality was tested using the FastQC script. Read files were imported to the Galaxy Web-based analysis portal ^47^. Reads were mapped to the human genome (hg19 assembly) using Bowtie2 ^48^. Properly paired CUT&RUN reads were filtered using a Phred score cutoff of 30 and mitochondrial reads were removed. Peaks were called with MACS2. To create profiles of CUT&RUN data, DeepTools 2 was used ^49^. Normalized bigWig files were created using bamcoverage with a bin size of 10 and normalizing to counts per million reads mapped (CPM). BigWig files were used to compute reads centered on YAP-seq peak summits (called with MACS2) using computeMatrix. Profiles were created with plotProfile tools. CUT&RUN data were visualized with the Integrated Genome Viewer ^50^. HISAT2 was used to align RNA-seq reads against hg19 ^51^. FeatureCounts and limma were used to analyze differential expression ^52–54^. DAVID was used for the identification of enriched GO terms. Gene Set Enrichment Analysis (GSEA) ^55^ was used to determine enriched gene sets.

## ACKNOWLEDGEMENTS

We thank Svenja Meierjohann for sharing reagents. We thank Susi Spahr for assistance with experiments. We thank Juan Pablo Prada Salcedo for help with analyzing tumor sections. This work was supported by grants from the Deutsche Krebshilfe (70112811) and Deutsche Forschungsgemeinschaft (DFG, GA 575/10-1) towards SG. B.v.E. was supported by grants from the BMBF (16GW0271K), DFG (EY 120/4-1), and the German Cancer Aid (Deutsche Krebshilfe; 70113138). The FLI is a member of the Leibniz Association and is financially supported by the Federal Government of Germany and the State of Thuringia. The animal facility as well as the core service histology of the FLI are gratefully acknowledged.

## AUTHOR CONTRIBUTION

SG planned the study; SG, BvE, DG, FL and MJ designed experiments; DG, FL, MJ, LB, FZ and VS conducted the experiments; SG, BvE, DG, FL and MJ analyzed data; SG performed bioinformatic analyses; CPA performed and supervised next generation sequencing; SG wrote the manuscript.

## DATA AVAILABILITY

RNA-sequencing and CUT&RUN datasets are available at the NCBI’s Gene Expression Omnibus ^56^ under the accession number GSE227974.

