## Supplemental Information for "Inhibition of the YAP-MMB interaction and targeting NEK2 as potential therapeutic strategies for YAP-driven cancers"

#### SUPPLEMENTAL FIGURE AND TABLE LEGENDS

**Supplemental Figure S1: MY-COMP inhibits YAP mediated transcription of MMB-target genes** A) Immunoblot analysis of cytoplasmic (C) and nuclear (N) lysates of MCF10A cells expressing ER-YAP2SA treated with 4-OHT for 14 hours or left untreated. Histone H2B served as a nuclear marker and tubulin as cytoplasmic marker. B) and C) RT-qPCR and immunoblot analyses of MCF10A ER-YAP2SA cells treated with 4-OHT for the indicated times (n=2 replicates). Actin served as a control for the immunoblot. D) MY-COMP was stably expressed in MCF10A-ER-YAP2SA cells. Cells were treated with or without doxycycline and the expression of MY-COMP was analyzed by immunostaining with an HA-antibody. Scale bar: 25 µm. E) and F) MCF10A cells stably expressing ER-YAP5SA and doxycycline-inducible MY-COMP were treated with 4-OHT and doxycycline as indicated. E) Immunoblotting was used to analyze the expression of the indicated proteins. Actin served as a control. F) RT-qPCR to analyze the expression of the *CYR61* mRNA. Error bars indicate s.d. of three biological replicates.

**Supplemental Figure S2: Expression of MY-COMP in melanoma cell lines.** The indicated uveal melanoma cell lines and cutaneous melanoma cell lines were stably transfected with doxycycline inducible MY-COMP. Dose dependent expression of MY-COMP constructs was verified by immunoblotting with an HA-antibody. Tubulin served as a control.

**Supplemental Figure S3: MMB-target genes regulation by MY-COMP and YAP**
GSE analysis of MMB-target gene signature upon MY-COMP expression or YAP/TAZ depletion in 92.1 cells.

**Supplemental Figure S4: shRNA mediated depletion of NEK2 inhibits the growth of uveal melanoma cell lines.** A) Schematic illustration of the lentiviral pInducer10 construct used to express the shRNA targeting NEK2 and turboRFP(tRFP). B) UM cell lines with inducible constructs for shRNAs against NEK2 or a luciferase (shctrl) as described in A) were generated. Cell lines were treated with 1µg/ml doxycycline for three days and *NEK2* expression relative to untreated control cells was analyzed by RT-qPCR. *GAPDH* expression was used for normalization. Error bars show SD of three independent experiments. C) NEK2 expression of cell lines described in B was analyzed by immunoblot. Vinculin served as a control. D) UM and CM cells were seeded at low density and left treated or untreated with the indicated concentrations of doxycycline for 10 days. Cells were fixed and stained with crystal violet. Plots show quantification of crystal violet staining. N=3 biological replicates each performed in triplicates. Error bars indicate SEM of one representative experiment.

**Supplemental Figure S5: Depletion of NEK2 in uveal melanoma cells results in apoptosis.** UM and CM cells were treated with the indicated concentrations of INH1 for 4 days. Cell cycle phases were determined by PI staining followed by FACS analysis. See Figure 6J for a quantification of cells in subG1.

**Supplemental Table 1**

List of primer sequences and siRNAs

**Supplemental Table 2**

List of antibodies

**Supplemental Table 3**

Significance for association with overall survival (OS) of uveal melanoma and skin cutaneous melanoma patients based on stratifying for expression quintiles of YAP/MMB target genes.

### Supplemental Figure S1

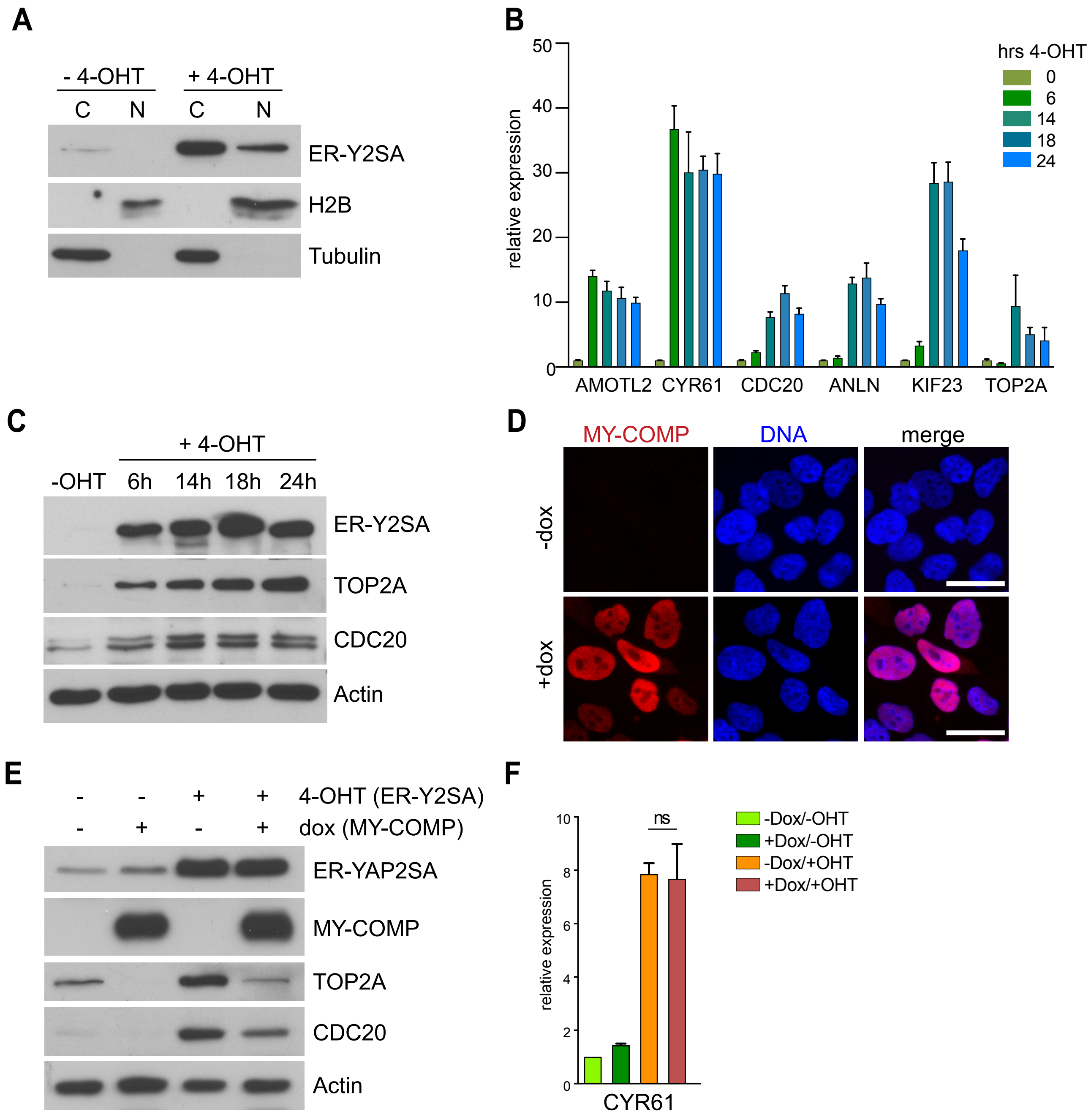

### Supplemental Figure S2

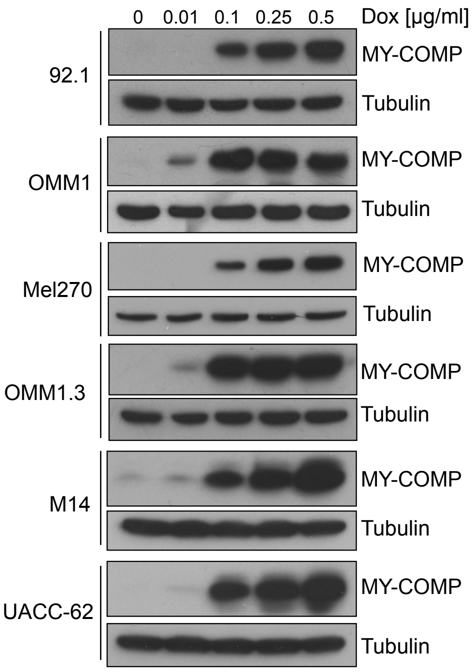

Supplemental Figure S3

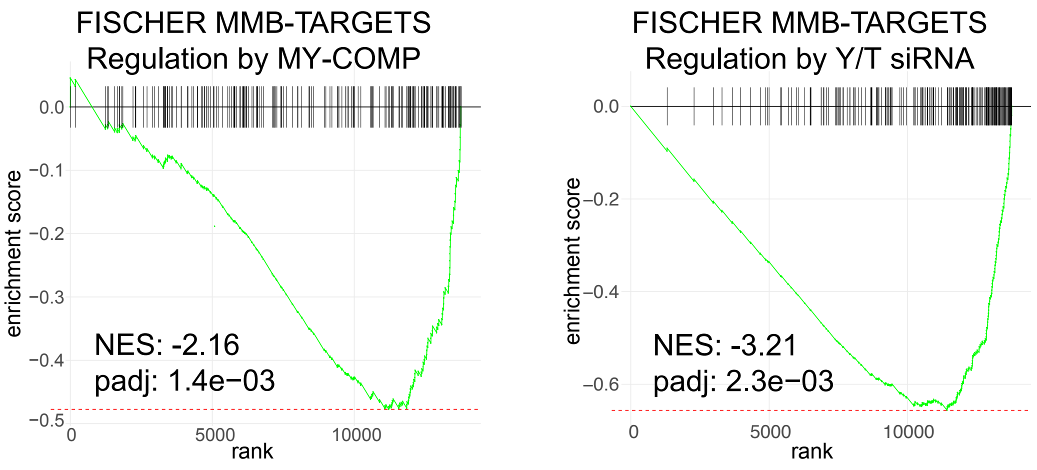

### Supplemental Figure S4

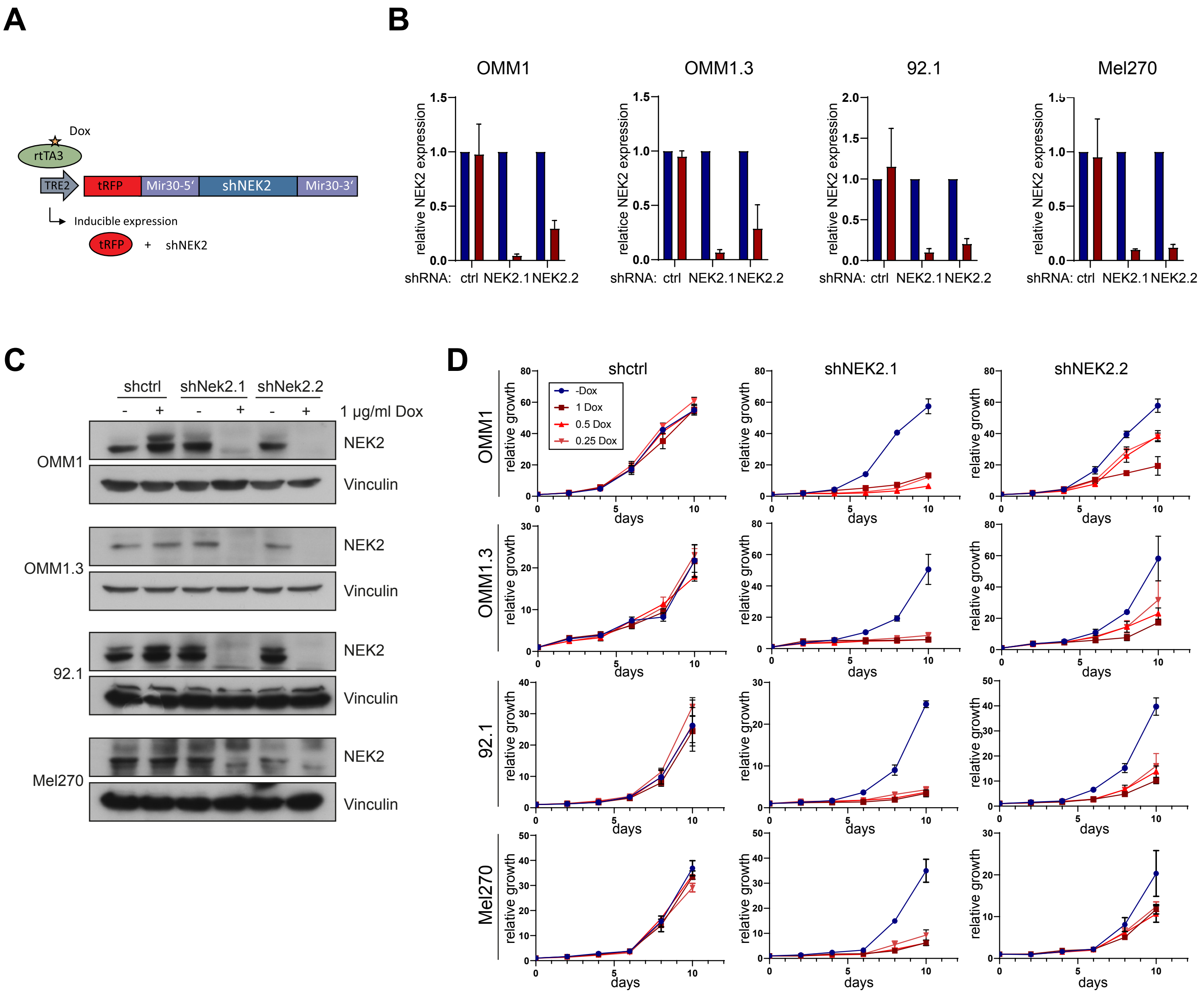

### Supplemental Figure S5

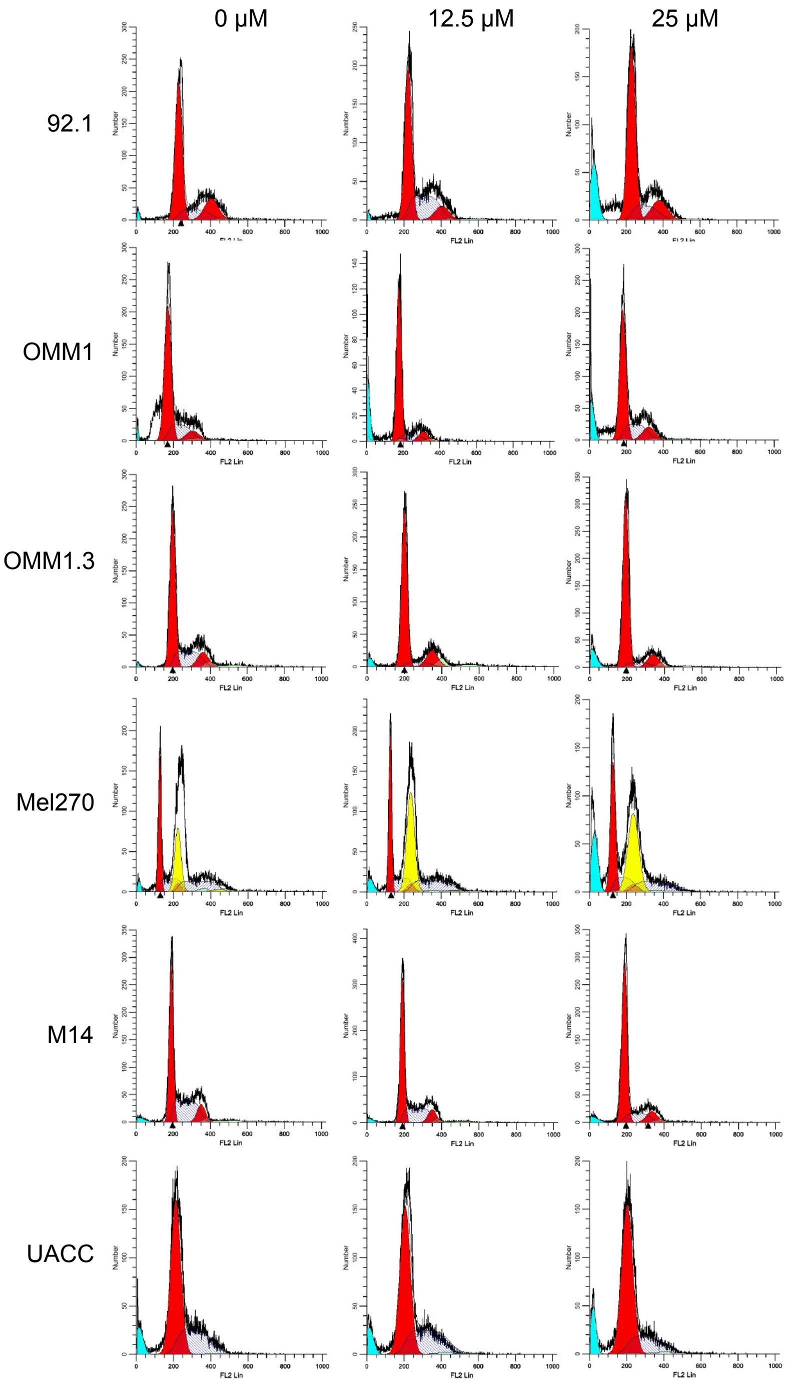

#### Supplemental Table 1: Oligonucleotide and siRNA sequences

##### RT-qPCR primers

| Gene | fw primer | bw primer |
|---|---|---|
| AMOTL1 | AGGCTGCAGAGAGACAATGAG | CTCAGAGAGCCGCTGGATT |
| ANLN | GCGAGCTAGACAGCCACTTT | TTTTTGATGGCGATGGTTTT |
| ARHGAP11A | CAGAACACCTTCTATTACACCTCAAG | GCATTTGGTGTAAGAATCACTGG |
| CYR61 | CAACCCTTTACAAGGCCAGA | TGGTCTTGCTGCATTTCTTG |
| DEPDC1 | CCTATGGAGAGTCAGGGTGTG | CGAAAAGATGTGGTAACTTCATTC |
| GAPDH | GCCCAATACGACCAAATCC | AGCCACATCGCTCAGACAC |
| KIF23 | CCTAACGTCCCGCAGTCTT | AGGTTTCCGGGGTGTCTTAG |
| KIF4A | TGGTCAGACAGCCCAGATG | TCTTCTAGCTTGGCGTTCATT |
| NCAPH | ACCTCAAACCAGGCACCA | TCTTCATAATGCTCAGTCTCTACCC |
| NEK2 | CATTGGCACAGGCTCCTAC | TCATGGAGCCATAGTCAAGTTCT |
| PLK1 | AAGATCTGGAGGTGAAAATAGGG | AGGAGTCCCACACAGGGTCT |
| TAZ | GTATCCCAGCCAAATCTCGT | TTCTGCTGGCTCAGGGTACT |
| TOP2A | TCTGGTCCTGAAGATGATGCT | TTAGTTAACCATTCCTTTCGATCA |
| YAP | GACATCTTCTGGTCAGAGATACTTCTT | GGGGCTGTGACGTTCATC |

##### 3C-qPCR

| Fragment number | sequence |
|---|---|
| anchor | GCCCTCTCAGGCACAAGAAAGG |
| 1 | GCCGTTTGTCATCTCTCACTGGG |
| 2 | GACTCTGGCCCTGAAGGAATGC |
| 3 | CCTAGAGCTGGGGCATTTGGC |
| 4 | GCTCAGGCAGTACTGCTGGTC |
| 5 | TACATGCCTGCAGCCCTCCTG |
| 6 | TGAGCCGAGATCATGCCACTGC |
| 7 | GGTTGTGGCACTGGAGACAGATTC |
| 8 | CGCGGAAGTGCAGAGAAGCAG |
| 9 | GGTCAAGCGCTGCAGAGATGC |
| 10 | GAGGCTTCAGGGCCATGATGAGG |
| 11 | GTCCCTAACCCCACACAGTCAGAC |
| 12 | CCCTCCAGGGTGACAAAGGGAC |
| loading control fw | GTCGCTTGAGCCCAGGAGTTC |
| loading control bw | CTCTTGGCCTCGAGCAATCCG |

##### siRNAs

| Gene | Name/ sequence | Reference |
|---|---|---|
| ctrl | Cat#4390843 | Thermo Fisher Scientific |
| YAP | UCUCUGACCAGAAGAUGUC | Azzolin Cell 2014; 158:157–70. |
| TAZ | ACGUUGACUUAGGAACUUU | Azzolin Cell 2014; 158:157–70. |
| B-MYB | GAAACGAGCCUGCCUUACAUU | Schmit Cell Cycle 2007; 6:15, 1903-1913 |
| LIN9 | GGAAGAGAGAUCAGCAUUAUU | Schmit Cell Cycle 2007; 6:15, 1903-1913 |

**Supplemental Table 2: Antibodies used in the study**

| Protein | host species | Source | Identifier | Application |
|---|---|---|---|---|
| alpha-tubulin | mouse | Sigma | T6074; RRID: AB_477582 | IF |
| anti-β-Actin | mouse | Santa Cruz Biotechnology | Cat# sc-47778; RRID: AB_626632 | WB |
| B-MYB | mouse | - | clone LX015.1; RRID: not available | WB |
| CDC20 | mouse | Santa Cruz Biotechnology | Cat# sc-13162; RRID: AB_628089 | WB |
| flag (M2) | mouse | Sigma | Cat# F3165; RRID: AB_259529 | WB |
| gamma-tubulin | rabbit | Sigma | T5192; RRID: AB_264690 | IF |
| HA (HA.11) | mouse | HISS | Cat# MMS-101P; RRID: AB_2314672 | WB |
| Histone H2B | rabbit | Abcam | Cat# ab1790, RRID:AB_302612 | WB |
| Histone H3 (acetyl K27) | rabbit | Merck | Cat# 07-360; RRID: AB_310550 | CUT&RUN |
| Histone H3 (mono methyl K4) | rabbit | Abcam | Cat# ab8895; RRID: AB_306847 | CUT&RUN |
| Histone H3 (tri methyl K4) | rabbit | Abcam | Cat# ab8580; RRID: AB_306649 | CUT&RUN |
| Histone H4 (acetyl K5,8,12,16) | rabbit | Abcam | #06-598 | CUT&RUN |
| IgG | rabbit | Sigma | Cat# I5006; RRID: AB_1163659 | CUT&RUN |
| IgG | mouse | Sigma | Cat#I5381;RRID:AB_1163670 | CUT&RUN |
| LIN9 | rabbit | Bethyl | Cat# A300-BL2981; RRID: N/A | WB |
| NEK2 | mouse | BD Biosciences | 610593; RRID: AB_397933 | WB |
| phospho-YAP (S127) | rabbit | Cell Signaling | Cat# 4911; RRID: AB_2218913 | WB |
| TOP2A | mouse | Santa Cruz Biotechnology | Cat# sc-365916; RRID: AB_10842059 | WB |
| Vinculin | mouse | Sigma | V9131; RRID: AB_477629 | WB |
| YAP | mouse | Santa Cruz Biotechnology | Cat# sc-101199;RRID: AB_1131430 | WB |
| YAP | rabbit | Cell Signaling | Cat# 14074; RRID: AB_2650491 | IHC |
| YAP | rabbit | Novus Biologicals | NB110-58358 ; RRID:AB_922796 | CUT&RUN |
| Anti-mouse HRP conjugated | | GE Healthcare | Cat# NXA931; RRID: AB_772209 | WB |

**Supplemental Table 3**

| gene | p-value UVM | p-value (SKCM) |
|---|---|---|
| NEDD9 | 0.000208753 | 0.037072884 |
| CDC25B | 0.000588943 | 0.441650859 |
| KIF20A | 0.001794255 | 0.033327048 |
| RRM2 | 0.005048059 | 0.411319142 |
| SKA1 | 0.009665771 | 0.163408569 |
| CDC25C | 0.010509242 | 0.078117992 |
| ASF1B | 0.010721858 | 0.044023866 |
| TPX2 | 0.022868364 | 0.270745245 |
| H2AFV | 0.027098992 | 0.116804935 |
| NEK2 | 0.028567756 | 0.650056761 |
| IQGAP3 | 0.031032975 | 0.001684824 |
| STIL | 0.035435754 | 0.727552757 |
| NUSAP1 | 0.038886372 | 0.98435199 |
| CKS1B | 0.046319613 | 0.17627508 |
| KIF18B | 0.050078456 | 0.169090775 |
| CHEK2 | 0.050956835 | 0.321538962 |
| AURKB | 0.055967001 | 0.006794848 |
| CCNA2 | 0.064948158 | 0.218099316 |
| RAD21 | 0.086082933 | 0.025857304 |
| CENPA | 0.101541651 | 0.031993769 |
| RACGAP1 | 0.114566307 | 0.452055672 |
| PIM1 | 0.121138151 | 0.040263734 |
| KIF2C | 0.148341343 | 0.012025081 |
| C15ORF23 | 0.199719692 | 0.61005831 |
| DBF4B | 0.241450035 | 0.06709812 |
| NCAPD2 | 0.250878195 | 0.203874428 |
| CENPN | 0.269690208 | 0.6310422 |
| FAM102B | 0.285514438 | 0.16497173 |
| INCENP | 0.329642012 | 0.098372102 |
| BRD8 | 0.419880382 | 0.098032762 |
| ESPL1 | 0.438853653 | 0.003614506 |
| DCAF16 | 0.628284186 | 0.375714613 |
| RAD54L | 0.828725955 | 0.147890486 |
